# Acute loss of fingertip sensation leads to general compensatory changes in eye-hand coordination

**DOI:** 10.1101/2023.11.28.569022

**Authors:** Kevin Ung, Jeffrey M. Yau, Per F Nordmark

## Abstract

The role of sensory feedback is well established in current models of motor control, evidenced by deficits in movement coordination resulting from impaired sensory function. When vision and touch are both available for object-oriented manual behaviors, these senses can be distinctly leveraged; vision guides movement planning while touch provides feedback on hand-object interactions. How eye-hand coordination changes with the loss of somatosensory feedback has not been well studied. Conceivably, vision is recruited to compensate for the feedback lost when touch is abolished. We tested healthy participants on a manual dexterity task, consisting of moving small metal pegs. The task was performed before and after administration of digital anesthesia that abolished cutaneous sensations while preserving motor function with the acting hand. During peg collection, transport, and placement epochs, we tracked gaze direction and hand positions while also recording forces applied to the pegboard. We hypothesized that the nervous system selectively adapts eye-hand coordination according to the dexterity demands of the task epochs. We found that participants maintained the ability to perform the pegboard task following the loss of cutaneous feedback, albeit with longer trial times and altered force profiles. Notably, somatosensory loss was accompanied by a shift in visual behavior marked by a closer alignment between gaze and hand positions during all task epochs, even those that did not involve object manipulation. Together, these data affirm the contributions of sensory feedback to force control in service of dexterous object manipulation and reveal the non-selective nature of compensatory eye-hand coordination processes.

**Significance Statement:** Touch and vision typically support distinct, but coordinated aspects of dexterous manual behaviors. Here, we evaluated how acute removal of tactile feedback using digital anesthesia affected performance and eye-hand coordination in a manual dexterity task. With insensate fingers and intact vision, participants continued to perform the task successfully, albeit with longer trial times and altered force profiles. We also observed closer alignment between gaze and hand positions during all task epochs, even those that did not involve object manipulation. Our results reveal the consequences of acute somatosensory loss and the general nature of compensatory eye-hand coordination processes.

## Introduction

Dexterous manual interactions with objects require tight coordination between sensory and motor systems. When all sensory channels are available, vision and touch support complementary roles: vision provides information that supports grasping and movement planning (Johansson et al., 2001), while touch provides direct feedback for performance monitoring and error correction (Johansson & Flanagan 2009). When sensory inputs are diminished or lost, the available input channels may be recommissioned to fill in critical information gaps.

Touch provides crucial information about hand state, grasped objects, and the physical interactions between the hand and grasped objects (Johansson & Flanagan 2009). Tactile signals from proprioceptors and cutaneous mechanoreceptors provide information about hand posture and movements. Tactile cues can also inform inferences about object properties such as shape (Jenmalm et al., 2003; Johnson & Hsiao, 1992), alignment of fingertips (Monzée et al., 2003), size and weight, and texture/friction (Bilaloglu et al., 2016; Johansson & Westling, 1984; Sathian et al., 1989). Critically, touch provides feedback about the physical interactions between the hand and the object signaling not just the force of the interactions (Westling & Johansson, 1987), but also friction and object slip events (Birznieks et al., 1998; Johansson & Westling, 1987). Accordingly, touch is critical for motor function in service of object manipulation. Given the tight coupling between the somatosensory and motor systems, impoverished or absent cutaneous sensations result in impaired manual dexterity and object manipulation deficiencies.

Vision supports our manual interactions with the environment in several ways and gaze behavior are a fundamental component to the planning and monitoring of visually guided actions (Johansson et al., 2001; Sailer et al., 2005). Vision supports object localization that guides reaching (Johansson & Flanagan, 2009) and supports inferences about shape, size, and weight to guide prehension (Gordon et al., 1993; Jenmalm et al., 2000; Jenmalm & Johansson, 1997; Lukos et al., 2007; Winges et al., 2003). In comparison to tactile feedback, vision only provides indirect information about the mechanical interactions between the hand and objects, such as the forces applied as well as the making and breaking of contact. What information is provided by vision varies in time according to context.

Effective sensorimotor behavior requires the association of actions and their sensory consequences. Individuals who have lost their somatosensory and proprioceptive feedback develop movement disorders (Chesler et al., 2016; Mahmud et al., 2017; Rao & Gordon, 2001; Yahya et al., 2019) and ultimately compensate for the loss of input through other mechanisms (Jenmalm & Johansson, 1997). Furthermore, individuals that have suffered from nerve injuries affecting fingertip sensation report higher demand on seeing their fingers when performing dexterous manipulations with objects, and have also been shown to have increased gray matter volumes in eye-hand coordination areas of the secondary visual cortex in the brain, suggesting increased dependence on vision to compensate impoverished somatosensation (Nordmark et al., 2018). Such compensatory changes may be considered a form of sensorimotor learning.Motor movements can sometimes be restricted to the context in which the movement occurred, based on the goal of the action and the sensory information available (Sober & Sabes, 2005). With hand-object interactions, visual processing that typically subserves reaching and grasping maybe be recruited to provide feedback to compensate for missing or impoverished somatosensory signaling. Whether this compensation is restricted to specific task states (i.e., evaluating whether an object is successfully grasped) or generalizes across contexts is unknown.

In this study, we asked how loss of tactile feedback in the fingers affects gaze behavior during a fine motor task testing manual dexterity and whether compensatory gaze changes are specific or generalized. To address these questions, we tracked hand movements and gaze direction as participants performed a pegboard test that required the collection, transport, and placement of small pegs. We assessed performance before and after we abolished tactile feedback through local anesthesia of the digits. We hypothesized that the loss of somatosensory feedback would induce compensatory gaze changes that were specific to only trial phases involving precise object manipulation. We found that removal of tactile feedback perturbed performance which visual compensation only partially rescued. Notably, compensatory gaze changes were not restricted to specific trial phases in the pegboard task and instead reflected a general yoking of gaze to the anesthetized hand.

## Materials and Methods

### Participants

Five healthy individuals participated in the study (mean age: 27.8; range: 26-30; 2 females). Four participants were right hand dominant (Edinburgh inventory, laterality index: 75-100) (Caplan & Mendoza, 2011) and 1 participant was left hand dominant (laterality index: −80). Four of the participants were right eye dominant whereas 1 was bilaterally dominant. All participants had normal or corrected-to-normal vision. All participants gave their written informed consent in accordance with the Declaration of Helsinki, and the Swedish Ethics Review Authority approved the study. No participants reported a history of nerve disorders or diabetes. All participants were paid for their participation (500 SEK).

### General overview

Participants performed a dexterous unimanual task comprising the collection, transport, and delivery of pegs (Figure 1A). A fixed forehead support defined the individual location of the head and eyes during performance of the task. Eye and hand positions were recorded throughout the performance of the task using recording devices mounted in fixed positions on a height-adjustable desk (Figure 1B). A monitor (480 x 270 mm) defined the depth of the workspace and displayed written instructions to the participant during the tasks. A 90-mm deep shelf with collection (outer) trays and the pegboard (middle) tray was mounted 165 mm from the top of the computer screen, serving as the basis for the workspace. Force transducers registered normal and torque forces produced by the participants’ unimanual actions on each of the trays. The defined workspace for the acting hand during the task was limited to an area approximately 40 cm in front and 10 cm below the eyes (for the left hand, from centered to approximately 20 cm left of a midpoint between the eyes, and respectively to the right for the right hand). Table height was adjusted to each participant enabling them to stand while performing the task. The workspace areas were defined to maximize comfort for the participant while minimizing occlusion of gaze tracking. Black fabric covered the tabletop and screens mounted on its back and sides to prevent potential distortion of gaze and hand tracking by reflective light. For the same reason the participants wore sleeves of black fabric covering their arms down to wrist level.

**Figure 1.**
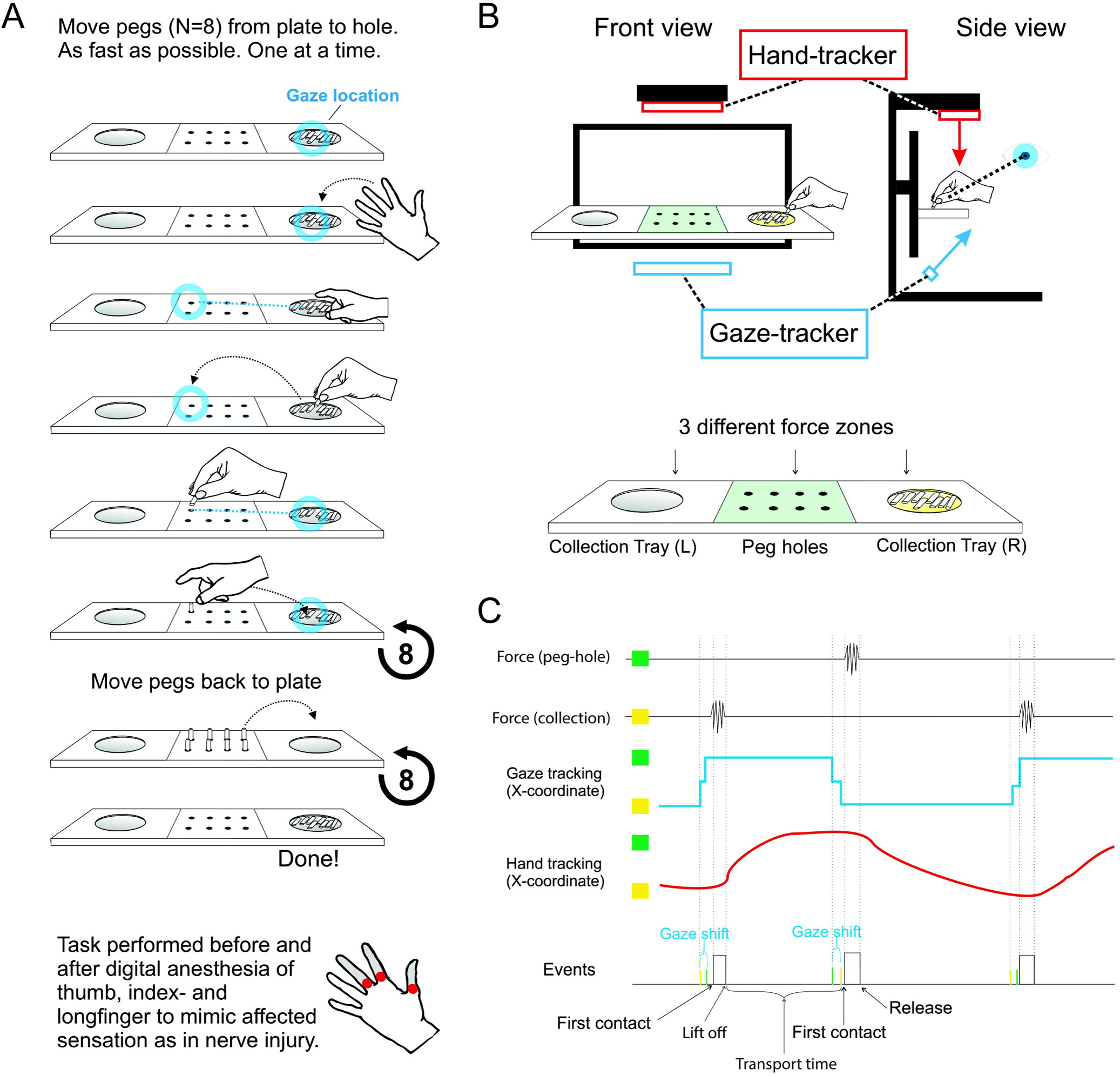
Overview of the workspace, task, and recorded data. (A) Schematic of the manual dexterity task. Participants pick up an individual peg from the collection tray and deliver the pegs to isolated holes in the delivery tray. This action is performed sequentially for all 8 pegs. Each tray has force transducers collecting normal and torque forces. (B) Layout of the positioning for the hand-tracker, gaze-tracker, and peg board. (C) Example traces of the force tracking, hand tracking, and eye tracking during the task.

### Pegs and peg-holes

Each peg consisted of a smooth metal rod (duralumin, diameter: 6 mm, length: 32 mm, weight: 7.6 g). All pegs had uniform color, shape, and weight. Peg-holes (diameter: 6.5 mm, depth: 10 mm) were arranged in 2 rows on a plastic 3D-printed surface (85 x 75 mm). Each row was separated by 23 mm and peg-holes within a row were separated by 13 mm. The front row was 65 mm from the computer monitor.

### Collection trays

The collection trays, one on the left and one on the right side of the peg-holes, were 3D-printed plastic concave discs with a 6-cm diameter. The ridge of the disc was aligned with that of the ledge, and the central part of the disc featured a flat surface with a 2-cm radius, situated 7 mm below the outer edge. We designed the tray in this way so that the participants were required to grasp the peg using a precision grip. Before the start of each block of trials, 8 pegs were placed flat in the collection tray such that no peg was on top of another peg.

### Gaze tracking

A gaze tracker (The EyeTribe tracker, Copenhagen, Denmark; sampling rate: 30Hz) was mounted centered below the workspace facing the eyes of the participant (Figure 1B). For each participant, before the start of the data registration for each round, the gaze tracker was calibrated using the device’s standard 9-point calibration procedure.

### Hand tracking

The device for tracking hand motion (Leap Motion controller, Mountain View, CA, USA; sampling rate: 120 Hz) was mounted centered 40-cm above the workspace (Figure 1B). Before the start of data registration, the device’s ability to identify the hand, thumb, index and middle finger coordinates were confirmed by the experimenter. Participants removed all jewelry and watches prior to the experiment to facilitate tracking.

### Force transducers

Force transducers (Load cell weight sensors - TZT HX711 Module Electronic Scale Aluminum Alloy Weighting Pressure Sensor, 1Kg, 1600 Hz, China) registered normal and torque forces applied to the peg-board tray and two collection trays. Continuous sampling from all devices enabled the registration of task-related force variations to eye and hand positions throughout the task (Figure 1C).

### Additional sensors

Light sensors were mounted in concave 3D-printed plastic hand rests (half spheres, radius: 3 cm). Participants placed their left and right hands in separate hand rests at the start of each block. The light sensors were used to confirm the movement onset and offset times for the active hand on each block. The light sensors also served to confirm the stationary status of the passive hand throughout the block.

### Procedure

At the start of each block, each hand was placed on top of hand rests equipped with light sensors. After a signal to start the task with the cued active hand, participants moved the pegs, one at a time, from the tray to the peg holes as fast as possible before returning the active hand to the hand rest. No instructions were given regarding how to pick up the pegs (other than one at a time) or regarding gaze direction (i.e., the participants were free to find their own strategy). If a peg was dropped, participants were instructed to continue by picking up a new peg and leaving the dropped peg for last. After a brief pause as both hands remained on the hand rests, another signal cued participants to retrieve the pegs from the peg-holes, one at a time, to return to the collection tray. After returning all the pegs from pegboard to the collection tray, participants placed the active hand back on the hand rest to signal the end of the block. With this design, a block comprised the placement of 8 pegs (placement trials) and the subsequent retrieval of 8 pegs (retrieval trials). For each trial, we identified the timing of collection, transport, and delivery events based on the force variations measured on the collection and peg-board trays (Figure 1C). Five blocks comprised a single round. Prior to digital anesthesia, participants performed 4 rounds consecutively with their dominant hand followed by 4 rounds with their non-dominant hand (160 placement and 160 retrieval trials with each hand). Digital anesthesia was then administered to their non-dominant hand (see below). After digital anesthesia, participants completed another 4 rounds with their dominant hand followed by 4 rounds with their anesthetized non-dominant hand. Participants took a short break between rounds. Gaze- and hand tracking calibration was performed prior to the start of each round. Across all rounds, each hand performed 320 total placement trials. With the non-dominant hand, 160 placement and 160 retrieval trials were performed with the effect of digital anesthesia. The entire study procedure was completed in approximately 3 hours.

### Digital anesthesia

Digital anesthesia was performed by a hand surgeon experienced in regional anesthesia in the hand and arm. To affect the fine motor control of the hand, we regionally anesthetized the three digits of the hand involved in dexterous manipulation: the thumb, the index finger, and the middle finger. To induce an immediate and persistent digital anesthesia effect we used a 1:1 mix of the fast onset local anesthetic Lidocaine (1%) and the long duration local anesthetic Levobupivacaine (0.5%). A 1.5-ml volume of this local anesthetic mix was injected in each of the addressed fingers with a volar approach through a 27 Gauge needle, with the participants in supine posture. This caused an ongoing loss of sensation of the addressed fingers and fingertips that persisted over the final 8 rounds of task performance (anesthetized period).

After the injection, but before the start of blocks with the anesthetized hand (20-45 minutes post-injection), sensory and motor functions of the anesthetized hand were evaluated using the two-point discrimination (2PD) test (measured in the radial and ulnar half of volar fingertip for thumb, index finger, and middle finger) and testing of range of motion (ROM) for the anesthetized fingers, respectively. All participants reported a loss of sensation for sharp touch and showed the expected sensory deficiency as indicated by dramatic 2PD threshold changes (15 mm or higher for all tested fingertips anesthetized compared to 3-4 mm pre-anesthesia). Importantly, there was no effect on ROM. There were no adverse effects, and all participants were able to complete the task after digital anesthesia.

### Data curation

We segmented the placement and retrieval trials into discrete collection, transport, and delivery epochs objectively using the normal force signals measured at the pegboard and collection trays. Collection epochs onset was defined as a change point as detected by the Primed Exact Linear Time (PELT) method (Killick et al., 2012) in the normal force (measured at the collection tray or pegboard) and collection epochs offset was defined by a change point with a return of normal force to baseline levels. Delivery epochs onset was defined as a change point in the normal force (measured at the pegboard or collection trays) detected by the PELT method and delivery epochs offset was defined by a change point with a return of normal force to baseline levels. Transport epochs were defined by each collection offset time and the subsequent delivery onset time. Given that the weight of pegs affected the measurements of the force transducers, each epoch trace was baseline subtracted so that the start of the event was normalized to 0.

### Data analysis

Analyses were performed using MATLAB (R2021a) and RStudio (R version 2022.07.01 Build 554). Our analyses consisted of 2 primary goals: 1) to determine the impact of digital anesthesia on peg collection, transport, and delivery, and 2) to determine how digital anesthesia modulated eye-hand relationship during peg collection, transport, delivery, and hand movement without a peg (i.e., from peg delivery to collection of the next peg). We further distinguished between trials involving peg collection from the large collection trays with delivery to the small peg-holes (precise delivery) and trials with collection from peg-holes with delivery to the large collection trays (coarse delivery). This distinction allowed us to test the hypothesis that eye-hand relationships depend on the task difficulty (i.e., precision demands).

To determine the impact of digital anesthesia on peg collection, transport, delivery, and hand movement without pegs, we quantified peg collection and delivery epoch durations (difference between onset and offset times), peak normal force, and total torque force. We also quantified peg transport durations (time elapsed from collection offset to delivery onset) and hand movement durations without pegs (time elapsed from delivery offset to collection onset). Separate analyses were performed for epochs extracted from the placement and retrieval trials. We report times rounded to the nearest hundredth of a millisecond and force to the nearest thousandth of a Newton or Newton-meter for normal and torque force, respectively. We defined the forces from the acting hand in downward direction against the tray/peg-holes as positive normal forces and computed absolute values to quantify normal force magnitude. To quantify torque force, we similarly computed the absolute value to estimate total force irrespective of direction along the recorded axis.

To determine how digital anesthesia modulated eye-hand relationships, we quantified the spatial separation between gaze location and index finger position during peg collection, transport, delivery epochs, and during peg-free hand movement. Given the configuration of the pegboard and collection trays, most of the positional variance during task performance was confined to the horizontal axis so we computed the distance between the gaze location and index finger (gaze-index separation index; GISI) in the x-axis. For collection and placement events, GISI was calculated during the time of peak normal force and as a function of normalized progress through the epochs defined as the percentage of the full epoch duration. GISI was similarly calculated as a function of normalized progress over the peg transport and peg-free hand movement events. Separate analyses were performed for epochs extracted from the placement and retrieval trials.

We statistically assessed how force, duration, and GISI were modulated according to hand and anesthesia state using generalized linear mixed-effects models (GLMM). To test modulation of force, duration, and GISI at peak force, we included anesthesia state (pre-anesthesia vs anesthetized) and hand (control hand vs anesthetized hand) as fixed effects. To account for subject-specific variance and testing time for each hand in the pre-anesthesia and anesthetized periods, we included the rounds nested within the participants as random effects. Thus, the GLMM took the form:

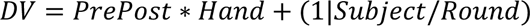

We similarly tested modulation of GISI as a function of normalized progress through the epochs by including percent progress through the epoch as an additional fixed effect. We used the lmerTest package (Kuznetsova et al., 2017) to provide tests of significance for all fixed effects, using the Satterthwaite method for degrees of freedom calculation. The reported degrees of freedom are rounded to the nearest integer.

## Results

We first present effects of digital anesthesia on performance during placement trials with respect to changes in force application and duration during the pick up and delivery epochs. We then present analogous results from the retrieval trials. Lastly, we present analyses of effects of anesthesia on eye-hand coordination during placement and retrieval epochs.

### Digital anesthesia effects on placement trial performance

To determine how digital anesthesia impacted performance during placement trials, we defined collection, transport, and delivery epochs using the force signals measured at the collection trays and pegboard. Figure 2 depicts the average force traces for individual participants and across the participant sample during coarse collection (i.e., peg pick up from trays). For the control hand, the normal force traces (Figure 2A) in the pre-anesthesia baseline condition (Control Pre-anesthesia: C_Pre_) and during anesthesia (Control During anesthesia: C_Dur_) rounds were highly similar. In contrast, the normal force traces for the anesthetized hand displayed obvious differences marked by greater forces which persisted longer in the anesthesized condition (Anesthetized During anesthesia: A_Dur_) compared to before anesthesia (Anesthetized Pre-anesthesia: A_Pre_). Visual inspection of torque force traces (Figure 2B) reveals the same distinction between the control and anesthetized hands.

**Figure 2.**
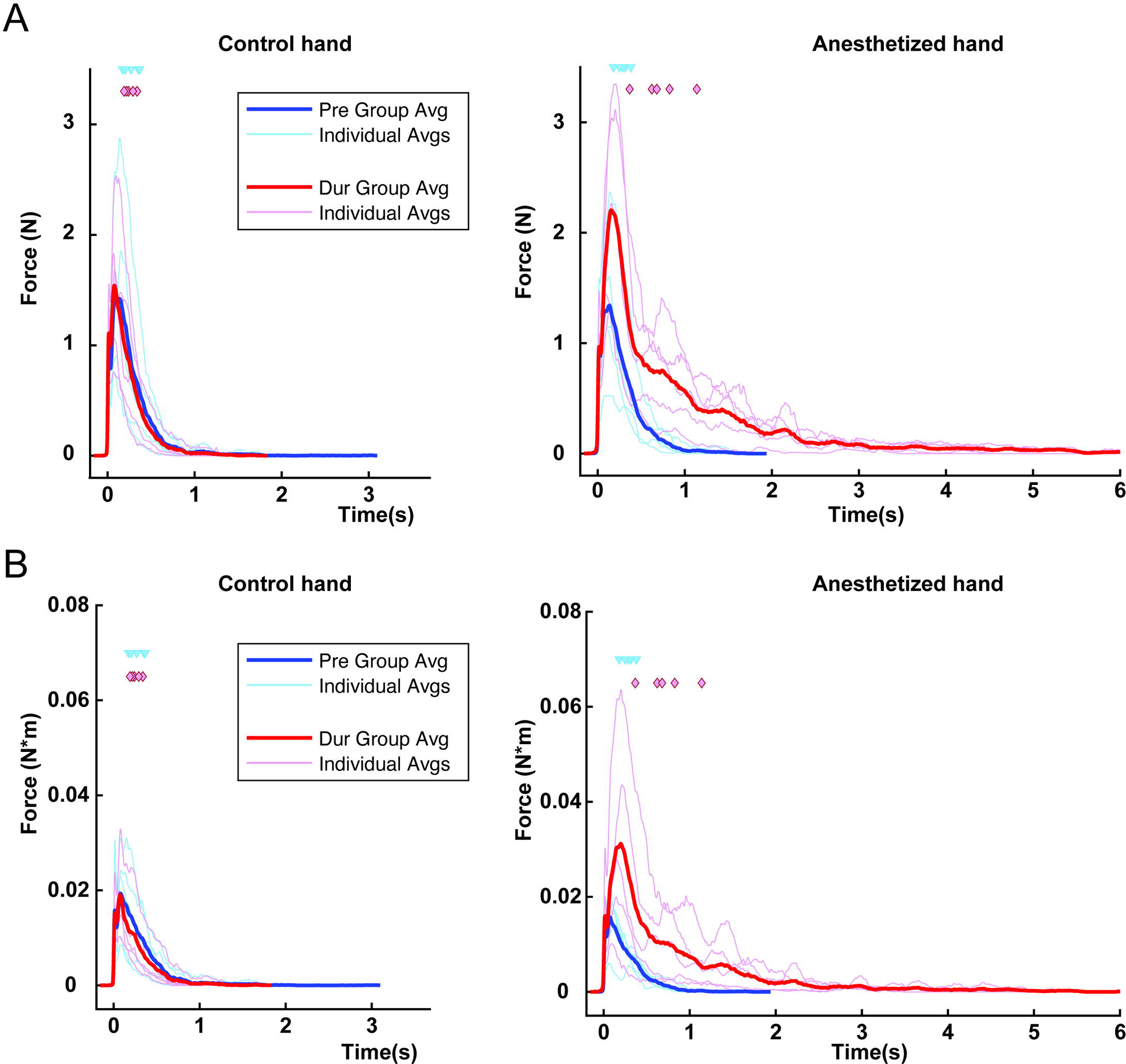
Duration of force application is increased during coarse collection with the loss of tactile feedback. (A) Traces of *normal force* produced during coarse collection of pegs during placement trials, i.e., trials with peg pick-up from tray, in the Control hand (left graph) and the Anesthetized hand (right graph) with onset of force aligned at *t* = 0. Light blue traces represent forces produced in trials before anesthesia per participant while dark blue trace represents the group average. Blue triangles represent the median duration of force application averaged across all trials per participant. Light red traces represent forces produced during trials with anesthesia, while dark red trace represents the group average. Red diamonds represent the median duration of force application averaged across all trials per participant. (B) Traces of *torque force* produced in the proximal/distal axis during coarse collection of pegs during placement trials in the Control hand (left) and the Anesthetized hand (right) with onset of force aligned at *t* = 0. Light blue traces represent forces produced before anesthesia per participant while dark blue trace represents the group average. Blue triangles represent the median duration of force application averaged across all trials per participant. Light red traces represent forces produced during anesthesia for each participant while dark red trace represents the group average. Red diamonds represent the median duration of force application averaged across all trials per participant.

To quantify collection performance, we computed the duration of normal force application by each hand before and after treatment with digital anesthesia to the non-dominant hand (Figure 3A). We predicted that somatosensory feedback loss would impair participants’ ability to collect pegs. Consistent with this prediction, force durations with the anesthetized hand (A_Dur_: 1132 ± 191.8 ms) were substantially longer compared to durations observed for the control hand during the pre-anesthesia and anesthesia rounds (C_Pre_: 348.7 ± 40.20ms vs C_Dur_: 317.2 ± 38.24ms) and the anesthetized hand prior to treatment (A_Pre_: 381.8 ± 21.43 ms). The GLMM (Table 3-1) captured this overall pattern as a significant treatment X hand interaction (*t*_3001_ = 16.696, P < 2e-16). Importantly, the GLMM failed to indicate significant main effects of treatment or hand. These results indicate that participants required more time to collect a peg successfully from the collection tray when anesthetized.

**Figure 3.**
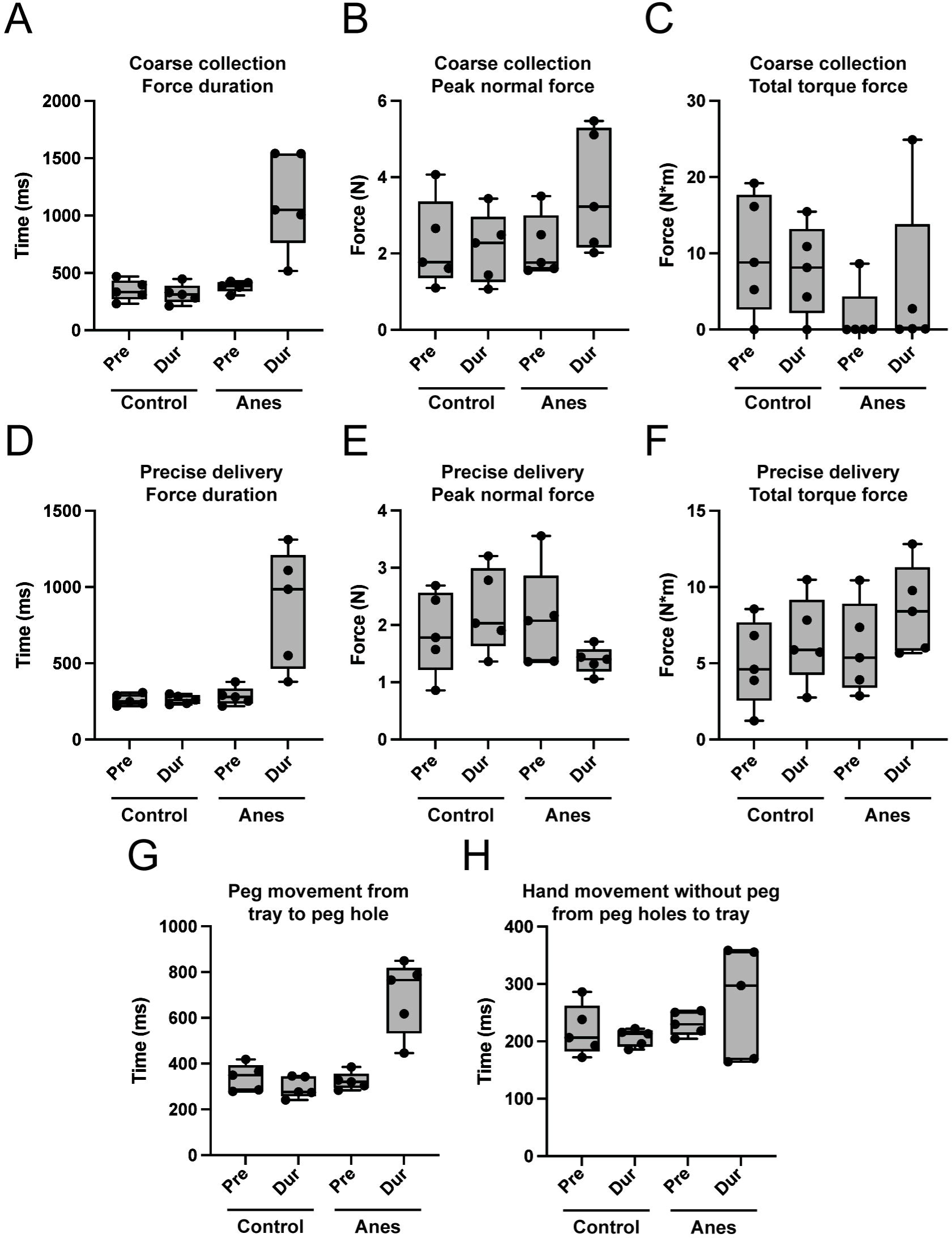
Force profiles during placement trials in the manual dexterity task. Boxplots of data from the placement epochs, i.e., peg pick-up from tray (coarse collection) and delivery to peg-holes (precise delivery). Coarse collection, box and whiskers (min to max) across all participants and conditions of (A) duration of force application in the collection tray, (B) peak normal force produced in the collection tray, and (C) total torque force in the collection tray. Precise delivery, box and whiskers (min to max) across all participants and conditions of (D) duration of force application in the peg-hole, (E) peak normal force produced in the peg-hole, and (F) total torque force in the peg-hole. Box and whiskers (min to max) depicting, for all trials sorted by condition, time elapsed between (G) completion of peg collection and initiating peg delivery for all trials sorted by condition, and (H) completion of peg delivery and initiating collection of the next peg.

Somatosensory feedback provides critical information regarding force production during object manipulation (Birznieks et al., 1998; Johansson & Westling, 1987; Westling & Johansson, 1987). Consistent with this notion, peak normal forces produced during collection using the anesthetized hand were more than 50% greater than forces produced pre-anesthesia and compared to control hand forces (C_Pre_: 2.242 ± 0.5215 N; C_Dur_: 2.142 ± 0.4163 N; A_Pre_: 2.185 ± 0.3699 N; A_Dur_: 3.626 ± 0.7119 N). The GLMM (Table 3-1) accounted for this pattern with a significant main effect of treatment (*t*_3002_ = −3.253, P = 0.001) and a significant treatment X hand interaction (*t*_3000_ = 20.162, P < 2e-16). The main effect of hand failed to achieve statistical significance (*t*_3001_ = −0.937, P =0.3489). Although there were qualitative differences in the total torque forces produced by the hands before and during anesthesia (Figure 3C), these were variable across participants and less suggestive of anesthesia effects. Indeed, the GLMM (Table 3-1) only revealed a significant main effect of hand (*t*_2463_ = −9.978, P < 2e-16) and failed to indicate significant main or interaction effects related to treatment. This pattern may reflect the use of different collection strategies with the dominant and non-dominant hands. Collectively, the force data imply that somatosensory feedback loss disrupts the regulation of normal forces, but not torque forces, during peg collection.

Digital anesthesia also impacted precise delivery of the pegs into the target peg-holes. The clearest evidence came from the force duration data (Figure 3D), which showed that participants required 3-fold more time to complete peg delivery when the hand was anesthetized compared to the other conditions (C_Pre_: 260.3 ± 16.56 ms; C_Dur_: 261.8 ± 13.33 ms; A_Pre_: 283.9 ± 26.73 ms; A_Dur_: 866.9 ± 174.3 ms). The GLMM (Table 3-2) accounted for this pattern with a significant treatment X hand interaction (*t*_3145_ = 10.449, P < 2e-16), but no significant main effects.

Digital anesthesia effects on peak normal force (Figure 3E) and total torque force (Figure 3F) measurements during peg delivery were less obvious and more variable across participants. Based on the GLMM (Table 3-2), peak normal force varied significantly according to hand (*t*_3144_ = 5.986, P = 2.40e-9) and time relative to anesthesia treatment (*t*_3143_ = 9.616, P < 2e-16). A significant treatment X hand interaction captured the increases in normal forces with the control hand and decreases with the anesthetized hand following treatment (*t*_3145_ = −19.472, P < 2e-16). These effects suggest hand-specific strategies that were modulated by digital anesthesia. One possibility is that participants rely on normal force feedback through touch to determine delivery success but shift to a strategy involving more lateral peg movements – which may be easier to appreciate by vision or intact somatosensory signals from more proximal regions – following somatosensory feedback loss. Indeed, torque forces increased in the rounds during anesthesia and were largest for the anesthetized hand (Figure 3F). This pattern was modeled (Table 3-2) as significant effects of time (*t*_1902_, P = 3.48e-3) and hand (*t*_1554_ = 1.981, P = 0.04) without a significant interaction effect. These collective results, combined with the duration data, reveal how precise peg delivery is impacted by digital anesthesia.

For completeness, we also assessed potential effects of digital anesthesia on peg transport and peg-free hand movement epochs, after participants had successfully collected or delivered the pegs, respectively. Peg transport durations were substantially longer in the anesthesia condition compared to other conditions (Figure 3G). This pattern was captured by a GLMM (Table 3-3) as a significant time X hand interaction (*t*_3002_ = 19.513, P <2e-16). The finding that peg transport took longer after somatosensory feedback loss implies that participants typically leveraged tactile signals during transport, likely to monitor stable grasping of the peg, and they may have shifted to an alternative monitoring strategy during digital anesthesia. Surprisingly, even hand movements between the pegboard and collection trays – after a peg had been delivered and before the collection of the next peg – were prolonged by digital anesthesia (Figure 3H). The GLMM (Table 3-3) also explained this variance with a significant time X hand interaction effect (*t*_2624_ = 2.387, P = 0.01). These results reveal the effects of digital anesthesia on task epochs that do not require dynamic interactions between the hand and pegs.

In sum, these analyses reveal the effect of digital anesthesia on collection, transport, and delivery events during placement trials. The modulation of event durations and forces support the conclusion that the loss of somatosensory feedback through digital anesthesia impacted pegboard task performance.

### Digital anesthesia effects on retrieval trial performance

Given the anesthesia effects observed during placement trials, we next looked for changes during retrieval trials (i.e., peg pick up from peg holes and delivery to collection tray). Compared to placement trial actions, peg retrieval requires a related, but different series of actions where there is a (precise) collection of an isolated peg (from a peg-hole) followed by transport of the peg for (coarse) delivery to the large collection tray. Participants repeated this action sequence until each of the 8 pegs were retrieved from the pegboard and delivered to the collection tray. We analyzed durations and forces during the collection, transport, and delivery events to determine digital anesthesia effects on retrieval trials.

With peg collection during the retrieval trials, we observed a clear increase in duration of force application in the anesthetized condition (Figure 4A). The GLMM (Table 4-1) accounted for this perturbation with a significant time X hand interaction (*t*_3152_ = 9.78, P < 2e-16). Digital anesthesia also systematically affected the forces recorded during precise collection: Maximum forces were recorded during collection with the non-dominant hand during anesthesia compared to the other conditions. The normal force patterns (Figure 4B) are consistent with the idea that somatosensory feedback loss disrupted force regulation during peg collection thereby increasing the forces generated during the dexterous manipulation event. The torque force patterns (Figure 4C) may reflect a general greater reliance on lateral peg movements (i.e., as a collection strategy) by the non-dominant hand that was accentuated by somatosensory feedback loss. Modulation of both normal and torque forces were captured by the GLMM (Table 4-1) as significant main effects of hand (normal: *t*_3151_ = 2.371, P = 0.02; torque: *t*_1612_ = 1.994, P = 0.04) and significant time X hand interactions (normal: *t*_3152_ = 7.286, P = 4.00e-13; torque: *t*_1815_ = 3.478, P = 5.17e-4). Collectively, these data reveal the influence of digital anesthesia on precise collection.

**Figure 4.**
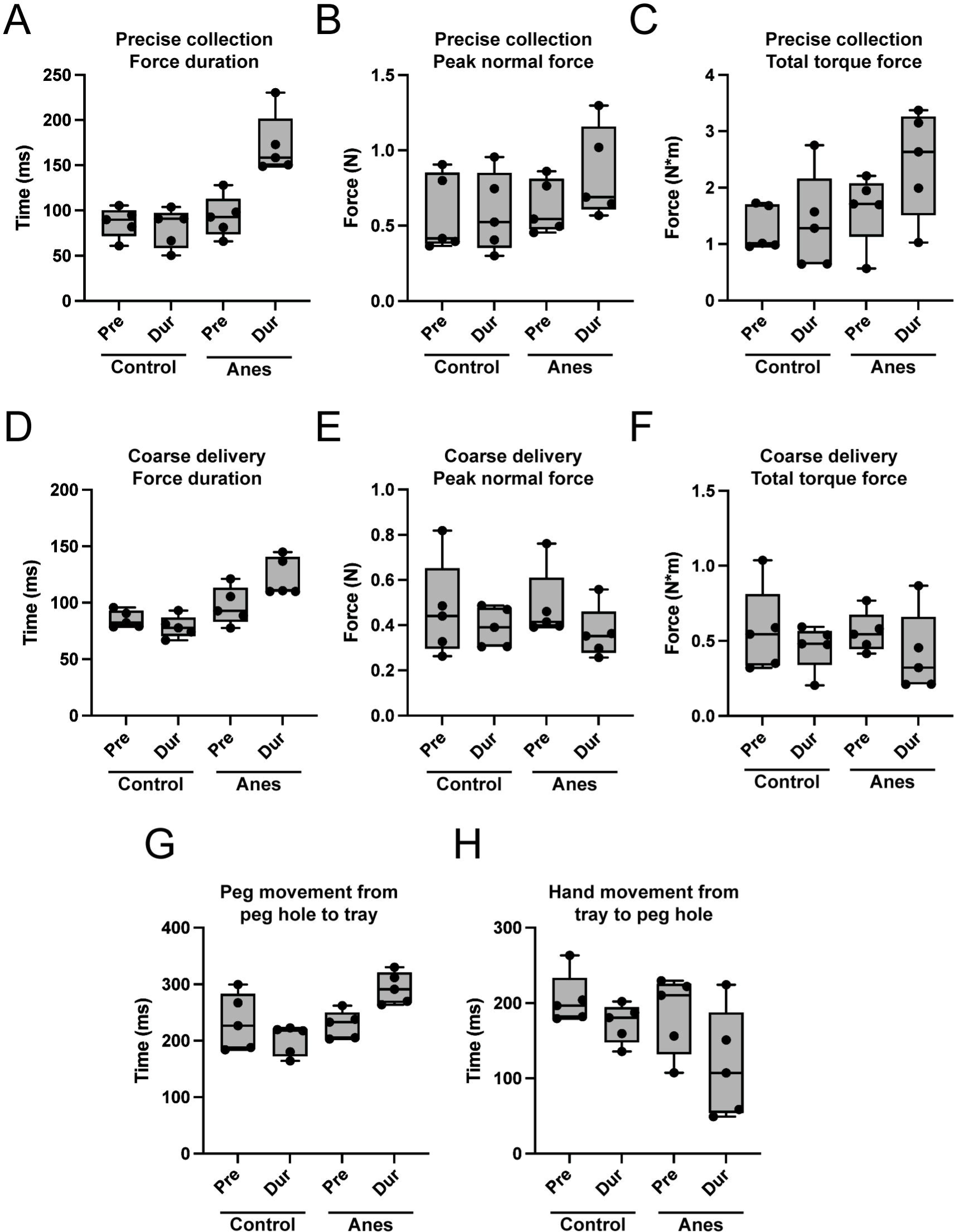
Force profiles during the retrieval trials in the manual dexterity task. Boxplots of data from the retrieval epochs, i.e., peg pick-up from peg-holes (precise collection) and delivery to tray (coarse delivery). Precise collection, box and whiskers (min to max) across all participants and conditions of (A) duration of force application in the peg-hole, (B) peak normal force produced in the peg-hole, and (C) total torque force in the peg-hole. Coarse delivery, box and whiskers (min to max) across all participants and conditions of (D) duration of force application in the collection tray, (E) peak normal force produced in the collection tray, and (F) total torque force in the collection tray. Box and whiskers (min to max) depicting, for all trials sorted by condition, time elapsed between (G) completion of peg collection and initiating peg delivery for all trials sorted by condition, and (H) completion of peg delivery and initiating collection of the next peg.

After peg collection from the pegboard, participants returned the pegs back to the collection tray. We predicted that digital anesthesia would have little impact on coarse delivery because the collection trays were relatively large, and we assumed delivery could be completed without a need for dexterous manipulation of the pegs on the tray. Surprisingly, normal forces persisted longer when coarse delivery was completed using the non-dominant hand with anesthesia compared to other conditions (Figure 4D). In fact, the GLMM (Table 4-2) revealed significant main effects of hand (*t*_3152_ = 3.96, P = 7.65e-5) and time (*t*_3153_ = −2.217, P = 0.03), as well as a significant time X hand interaction (*t*_3153_ = 7.313, P = 3.29e-13). To identify additional evidence for digital anesthesia effects on coarse delivery, we assessed the forces measured on the collection trays. While we found that normal forces (Figure 4E) and torque forces (Figure 4F) were reduced in the rounds completed during anesthesia, these reductions occurred with both the control and anesthetized hands which argues against specific digital anesthesia effects. Indeed, the force patterns were modeled (Table 4-2) as significant main effects of time (normal: *t*_3152_ = −4.876, P = 1.14e-6; torque: *t*_2502_ = −4.54, P = 5.90e-6) with no significant main or interaction hand effects. Thus, only the force duration data offered evidence that digital anesthesia impacted coarse delivery.

Based on our results showing that somatosensory feedback loss modulated peg transport and peg-free hand movement durations during placement trials, we predicted similar anesthesia effects during retrieval trials. Consistent with our predictions, transport time from peg collection to peg delivery increased in the anesthesia condition compared to pre-anesthesia and control hand performance (Figure 4G). The GLMM (Table 4-3) accounted for this pattern as a significant time X hand interaction effect (*t*_3138_ = 9.394, P < 2e-16). In contrast to the anesthesia effects on peg transport times, somatosensory feedback loss did not obviously impact hand movement times measured between peg delivery and the start of the next collection event (Figure 4H). Because model fitting failed to converge on a unique solution, we could not identify significant effects of hand or time on these peg-free hand movement times. Thus, during placement trials, somatosensory feedback loss only clearly modulated peg transport times. Together, these analyses reveal the effect of digital anesthesia on collection, transport, and delivery events during retrieval trials. Thus, the loss of somatosensory feedback through digital anesthesia impacted performance in all phases of both the placement and retrieval trials of the pegboard task.

### Eye-hand coordination compensation in placement trials following digital anesthesia

Having demonstrated that digital anesthesia modulated performance times and forces generated in the pegboard task, we next explored how somatosensory feedback loss impacted the spatio-temporal relationship between participants’ gaze direction and hand location during the task. When grasping and transporting objects with intact senses, vision serves primarily to guide planning of subsequent hand movements (Johansson et al., 2001), so gaze position typically leads hand position in space. In this mode, somatosensory signals provide feedback regarding the physical interactions between the hand and grasped object (Johansson & Flanagan 2009). In the context of our pegboard task, somatosensory feedback would signal the forces applied during peg collection, successful peg grasping and delivery, and stable peg handling during transport. With somatosensory feedback loss, we hypothesized that vision would be recruited to provide substitute feedback about the interactions between the hand and grasped object. Accordingly, we predicted that gaze position would be more closely tied to hand position after digital anesthesia.

We first examined eye-hand relationships by analyzing all peg-free hand movement epochs leading up to coarse collection (i.e., hand movements after peg delivery to the peg holes to peg pick up from the collection tray) (Figure 5A). Given that nothing was held in the hand during this epoch, we did not predict eye-hand differences related to anesthesia for these events. Contrary to our prediction, we observed that participants’ gaze position was significantly closer to the index finger (gaze-index separation index; GISI, see Methods) following digital anesthesia compared to the other conditions (intercept; hand X treatment, *t*_13200_ = −8.719; P < 2e-16) (Table 5-1). We next considered how GISI evolved over the movement period leading up to coarse collection. Although GISI changed under each condition, with spatial separation peaking 25-50% in the epoch, a GLMM revealed no significant treatment X hand X progress interaction (slope; Table 5-1). These results indicate that the reduced eye-hand separation in the during anesthesia condition compared to other conditions was maintained throughout the hand movement epoch. The fact that gaze was more closely anchored to the hand during peg-free hand movements suggests a compensatory mechanism that does not require the hand to be grasping an object.

**Figure 5.**
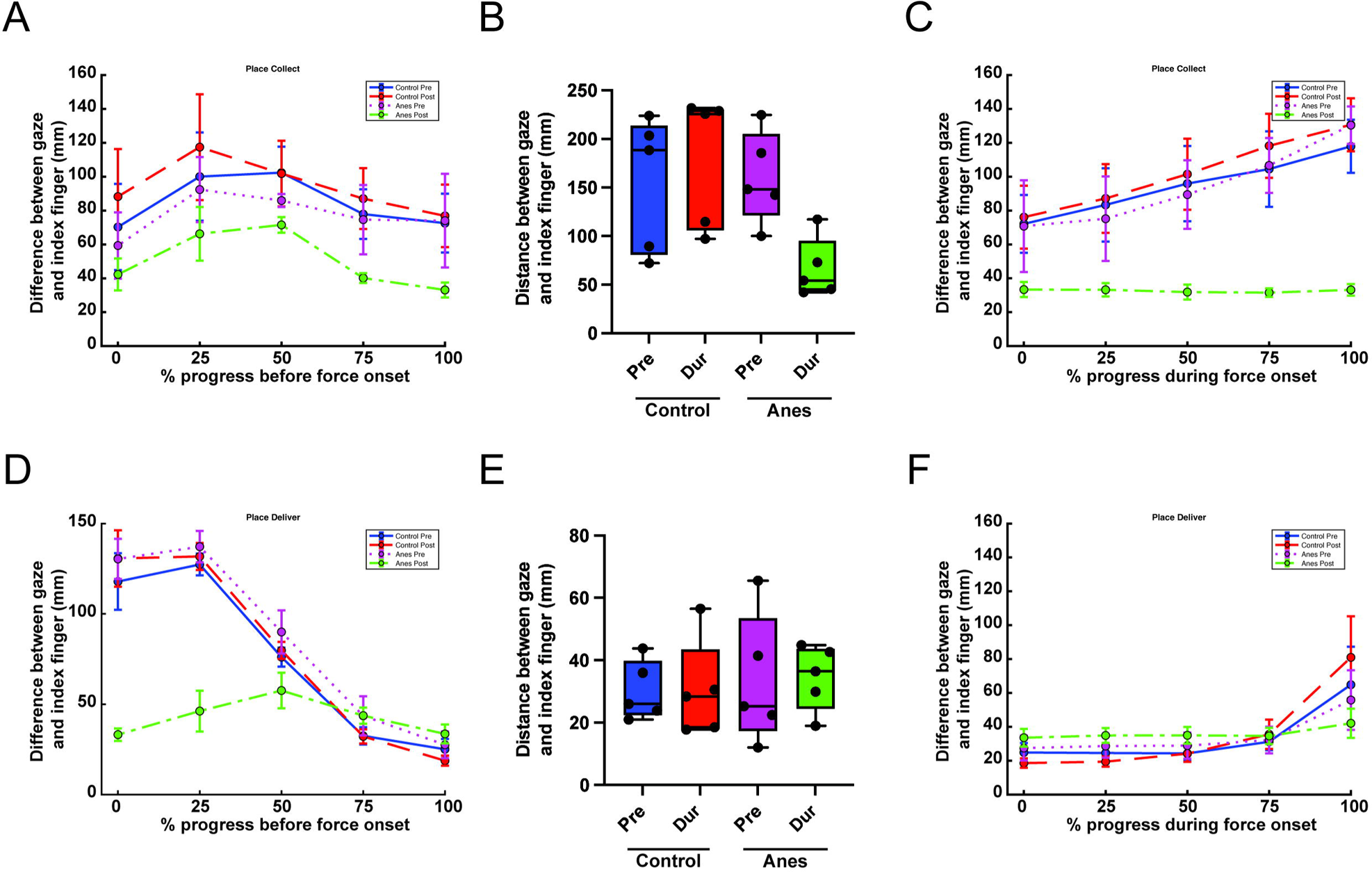
Participants maintained a closer gaze/hand relationship during the placement trials. (A) Line plot representing the difference between GISI through the transport time leading up to coarse collection across conditions, i.e., during peg-free hand movements. Participants’ eyes followed their hands closer after local anesthesia, although the relative distance compared to controls did not change through progress of the epoch. Data are represented as mean ± s.e.m. (B) Box and whiskers (min to max) showing the difference in spatial relationship between gaze and hand in all trials separated by condition at the event of peak normal force during coarse collection. At peak force application, GISI was consistently closer in anesthetized hands. (C) Line plots representing the difference between GISI through the progression of coarse collection epochs across conditions. Participants maintained a consistent gaze/hand relationship when the anesthetized hand was used. However, in all conditions with normal sensation, participants began to look ahead during the coarse collection epoch or even before the start of the epoch, as suggested by an increase in the gaze/hand relationship and thus a positive slope and a difference in GISI at 0 % progress of collection. Data are represented as mean ± s.e.m. (D) Line plots representing the difference between GISI through the transport time leading up to precise delivery at peg-holes across conditions. Participants initially have a large gaze/hand relationship with the gaze at peg-holes. As the epoch progresses towards targeted delivery, the gaze/hand relationship narrows as the hand holding the peg approaches the peg-hole. Under local anesthesia, gaze/hand relationship remains relatively consistent, equal to control levels at the end of the epoch when visual feedback is most important for reaching the target of putting down peg in peg-hole. Data are represented as mean ± s.e.m. (E) Box and whiskers (min to max) showing the difference in spatial relationship between gaze and hand in all trials separated by condition during peak force of precise delivery. At peak force application, we observed no differences in GISI across all conditions. (F) Line plots representing the difference between GISI through the progression of precise delivery across conditions. Visual feedback is highly important as can be seen in the closer gaze/hand relationship. Towards the last quartile of this epoch, the spatial distance between gaze and hand begins to increase in all conditions except post local anesthesia. Data are represented as mean ± s.e.m.

We next quantified the eye-hand relationship during coarse collection. We first examined GISI at the time of peak normal force production (Figure 5B), reasoning that participants were maximally engaged with the pegs at this time. Indeed, at this time point participants consistently had a lower GISI with anesthesia compared to other conditions as shown by a GLMM with a significant hand X treatment interaction (*t*_3001_ = −21.137, P < 2e-16) (Table 5-2). Because GISI is closer at peak normal force, we predicted that this may extend to the rest of the collection epoch. Given that the time of peak normal force varies across trials throughout coarse collection, we next modeled GISI as a function of coarse collection progression (Table 5-3). At the onset of the collection epoch, we observed lower GISI values in the anesthetized condition, a direct carry-over from the hand movement epoch (intercept; hand X treatment, *t*_15090_ = −7.617, P = 2.76e-14) (Figure 5C). GISI increased monotonically over the epoch under all conditions except in the anesthetized condition where gaze was generally fixed on the hand (Figure 5C). The GLMM captured this pattern as a significant hand X treatment X progress interaction (slope; *t*_15090_ = −12.6, P < 2e-16). These data suggest that participants were typically less reliant on gaze to accomplish coarse collection and quicker to look ahead to the next trial phase in control conditions. In contrast, participants looked closer at their hands and maintained this eye-hand relationship during coarse collection following the loss of somatosensory feedback.

For precise delivery, participants held the peg throughout transport to the pegboard. At the start of this epoch (Figure 5D), in continuity with the end of coarse collection, participants maintained a closer relationship between their gaze and hand positions in the anesthetized condition as compared to the other conditions (intercept; hand X treatment, *t*_15090_ = −42.084, P < 2e-16) (Table 5-4). In the anesthetized condition, GISI remained relatively low throughout the epoch. In contrast, in conditions with intact somatosensation, the relatively large GISI values at the start of the epoch declined monotonically over the transport period. A GLMM accounts for these patterns with a significant hand X treatment X progress interaction (slope; *t*_15090_ = 31.22, P < 2e-16). Together, these results support our hypothesis that vision, which typically precedes and guides hand movements, is recruited to monitor the status of the grasped peg during transport following somatosensory feedback loss.

Lastly, we examined eye-hand relationships during precise delivery when participants were tasked with placing the peg into a small hole on the pegboard (Figure 5E, F). We reasoned that this action requires more precision and a higher degree of eye-hand coordination compared to the other task phases even with intact somatosensation. Accordingly, we did not predict substantial differences in GISI values across conditions during the precise delivery epoch. Consistent with our prediction, GISI values were comparable across all conditions (hand X treatment, *t*_3144_ = 0.538, P = 0.59) at the time of peak normal force production (Figure 5E, Table 5-2). Visual inspection of GISI dynamics over the delivery epoch revealed modest across-condition differences and minimal across-time variation over most of event interval (Figure 5F). Notably, GISI increased sharply during the last event quartile in all conditions except for the anesthetized hand. These time courses may reflect a process in which gaze is initially required to perform precise delivery but is more quickly released – in conditions with reliable somatosensation – to prepare for the next trial phase. The GLMM accounted for these dynamics with a significant hand X treatment interaction on the intercept estimates (*t*_15810_ = 7.258, P = 4.11e-13) and a significant hand X treatment X progress interaction on the slope estimates (*t*_15810_ = −9.211, P < 2e-16) (Table 5-5). Together, these results indicate modest differences in eye-hand relationships during precise delivery, an action that generally requires greater involvement of vision. Importantly, GISI patterns late in the epoch support that notion that the loss of somatosensory feedback enhances participants’ reliance on vision to complete precise peg delivery.

In sum, these analyses revealed the effect of digital anesthesia on eye-hand relationships during collection, transport, and delivery events during placement trials: Gaze tracked more closely with the hand following the loss of somatosensory feedback. GISI changes related to digital anesthesia were obvious during coarse collection and peg transport events, implying that vision was recruited to support feedback mechanisms typically governed by touch. GISI modulation during precise delivery may be more subtle because this action typically requires more visual support. Notably, the closer eye-hand relationship during peg-free hand movements suggests a general compensatory mechanism in which vision becomes spatially yoked to the hand even in the absence of object manipulation.

### Eye-hand compensation in retrieval trials following digital anesthesia

We next examined eye-hand relationships during retrieval trials comprising precise collection, transport, and coarse delivery. Analysis of the peg-free hand movement epochs (preceding precise collection) additionally enabled us to evaluate whether eye-hand compensatory changes were specific to actions involving object manipulation.

The peg-free hand movement epochs in retrieval trials were generally characterized by an initial large eye-hand separation that decreased monotonically over the movement epoch (Figure 6A). This pattern is consistent with the notion that precise collection is a visually guided process in which participants initially directed their gaze to the pegboard to localize the target peg before they subsequently moved their hand to grasp the peg. Notably, GISI values across the movement epoch were significantly lower in the anesthetized condition compared to control conditions (intercept; hand X treatment interaction, *t*_13800_ = −13.129, P < 2e-16) (Table 6-1). GISI values reduced at different rates for the control and anesthetized conditions, and these differences were captured by a significant hand X treatment X progress interaction (*t*_13800_ = 9.471, P <2e-16) on slope estimates in the GLMM. These results indicate that participants directed their gaze closer to their hands following the loss of somatosensory feedback during hand movements preceding precise collection, consistent with the results in the peg-free hand movement epoch during coarse collection. These results are inconsistent with the hypothesis that eye-hand compensation is event specific and dependent on interactions with objects.

**Figure 6.**
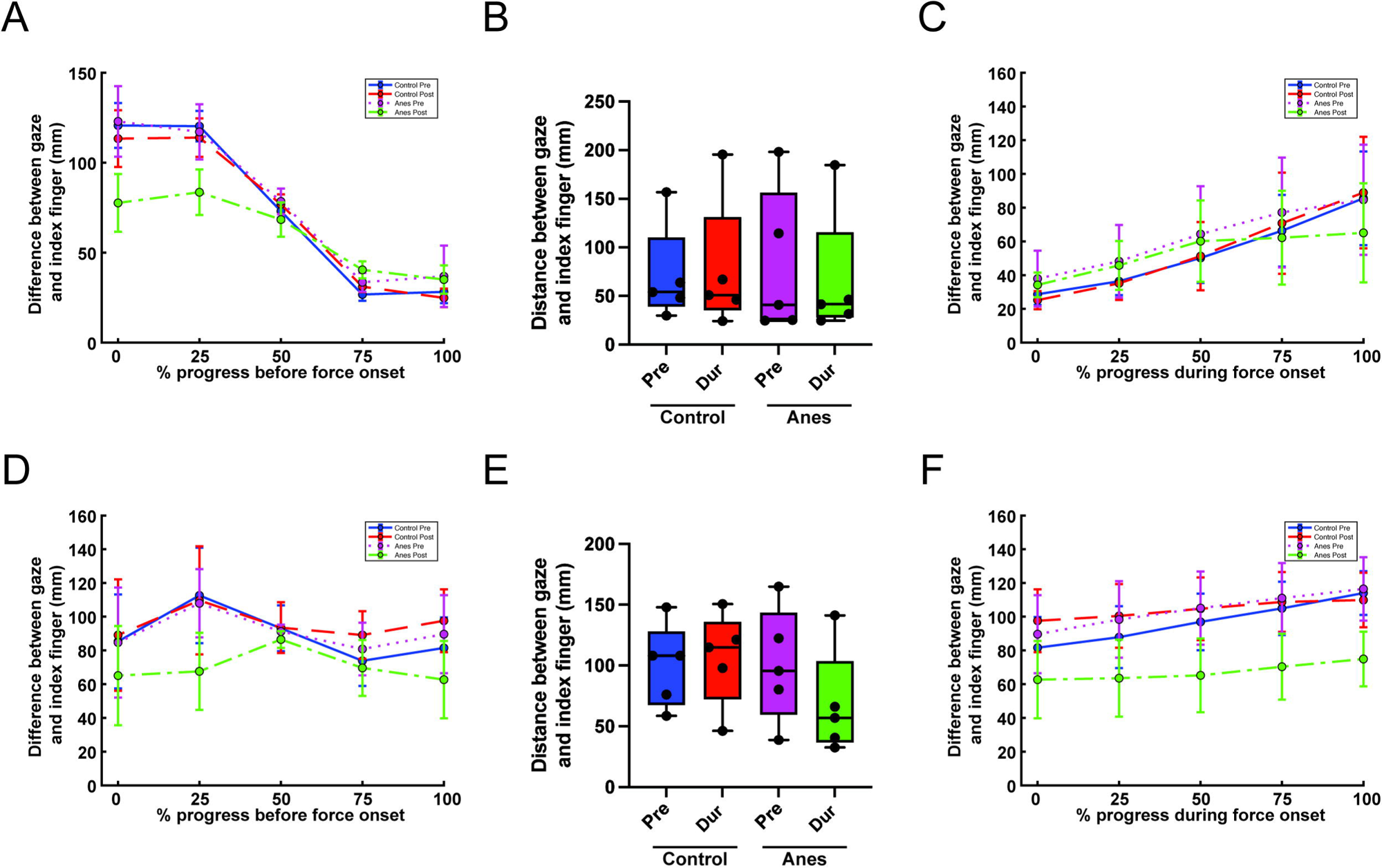
Participants maintained a closer gaze/hand relationship during specific epochs of the retrieval trials. (A) Line plots representing the difference between GISI through the transport time leading up to precise collection across all conditions. Participants had a comparable gaze/hand relationship pattern to that of precise delivery under control conditions. Under local anesthesia, participants had a closer initial gaze/hand relationship before converging with controls as the epoch progresses. Data are represented as mean ± s.e.m. (B) Box and whiskers (min to max) showing the difference in spatial relationship between gaze and hand in all trials separated by condition during peak force of precise collection. At peak force application, GISI was slightly lower after local anesthesia as compared to controls. (C) Line plots representing the difference between GISI through the progression of precise collection across conditions. Gaze/hand relationship increases in all conditions, suggesting a shift in gaze away from the peg and towards the target. In anesthetized conditions, the slope plateaus after 50% progression, an indication that there is still an increased reliance on vision as compared to controls. Data are represented as mean ± s.e.m. (D) Line plots representing the difference between GISI through the transport time leading up to coarse delivery across conditions. Participants begin with a smaller gaze/hand relationship in the anesthetized condition compared to controls. Through progression of the transport to the outer tray, we observed no significant change in slope between conditions. Data are represented as mean ± s.e.m. (E) Box and whiskers (min to max) showing the difference in spatial relationship between gaze and hand in all trials separated by condition during peak force of coarse delivery. At peak force application, GISI was consistently closer after local anesthesia. (F) Line plots representing the difference between GISI through the progression of coarse delivery across conditions. We again observed a closer gaze/hand relationship throughout the epoch in the anesthetized condition with no significant differences in slope. Data are represented as mean ± s.e.m.

To quantify eye-hand relationships during precise collection, we first evaluated GISI values when participants produced peak forces on the pegboard (Figure 6B). A GLMM captured a modest reduction in GISI in the anesthetized hand condition compared to other conditions with a hand X treatment interaction (*t*_3151_ = −3.172, P = 1.53e-3) (Table 5-2). Critically, mean GISI values during peak force production under all conditions were comparable to the mean GISI (at peak force) measured with the anesthetized hand during coarse collection (Figure 5B), conceivably reflecting the level of gaze demands that are intrinsic to precise collection. Across the precise collection epoch (Figure 6C), GISI values were similar across all conditions (intercept; hand X treatment, *t*_15850_ = 0.843, P = 0.399) (Table 6-2). GISI under each condition also exhibited parallel increases over event progression until diverging for the last quartile in the anesthetized condition. The GLMM accounted for this pattern with a significant hand X treatment X progress interaction on slope estimates (*t*_15850_ = −4.599, P = 4.27e-6). Importantly, the distances separating gaze and hand positions were substantially lower than those measured in the control conditions during coarse collection (Figure 5C), which were also marked by monotonic increases. Together with the precise collection analyses, the across-event comparison suggests that precise collection impose a greater gaze demand compared to coarse collection which may only be subtly enhanced by the loss of somatosensory feedback.

We next evaluated anesthesia effects on eye-hand relationships during peg transport preceding coarse delivery (Figure 6D) (i.e., peg pick up from peg-holes to delivery to collection tray). Given our hypothesis that vision is recruited to provide object manipulation feedback when touch is lost, we predicted that gaze would be more closely tied to the hand during peg transport with digital anesthesia compared to conditions with intact touch. Consistent with this prediction, we observed significantly reduced GISI values in the anesthetized condition (intercept estimates; hand X treatment interaction, *t*_15770_ = −5.686, P = 1.33e-8) (Table 6-3). Despite these magnitude differences, GISI values varied over the transport epoch similarly in all conditions (Figure 6D) and the similarity of the temporal profiles was confirmed by the non-significant hand X treatment X progress interaction on GLMM slope estimates (*t*_15770_ = −1.656, P = 0.097). These collective results indicate a tighter eye-hand relationship following somatosensory feedback loss that was generally maintained over the duration of the peg transport epoch.

Lastly, we compared GISI values during coarse delivery. We first quantified GISI values during peak force production (Figure 6E, Table 5-2) and found a significant reduction in the anesthetized hand condition compared to other conditions (hand X treatment interaction, *t*_3158_ = −9.223, P < 2e-16). Temporal analysis revealed that the tighter eye-hand relationship was evident over the entire coarse delivery epoch (Figure 6F). A GLMM captured the temporal patterns as a significant hand X treatment interaction on intercept estimates (*t*_15880_ = −13.823, P < 2e-16) and a non-significant hand X treatment X progress interaction on slope estimates (*t*_15880_ = 0.94, P = 0.35) (Table 6-4). Taken together, these results reveal that the loss of somatosensory feedback is associated with a tighter eye-hand relationship that is generally consistent across the entire coarse delivery epoch, again implying that vision was recruited to compensate for the loss of feedback mechanisms typically governed by touch.

## Discussion

In this study, we examined eye-hand coordination during a dexterous motor task when fingertip somatosensation was acutely abolished. Participants performed a unimanual pegboard task before and after digital anesthesia was administered to the thumb, index finger, and middle finger on one of the hands. We separated the task into placement and retrieval trials, each consisting of four specific epochs: peg collection, transport, and delivery, along with peg-free hand movements. Because pegs were collected from and delivered to large and small holding containers, we examined whether somatosensory feedback loss impacted task performance in a manner that depended on specific actions according to their required precision. With insensate fingers, participants continued to complete the task successfully, albeit with longer trial times and altered force profiles. When examining the gaze-hand spatial relationship throughout the task, we found that gaze was generally directed closer to the hand under anesthesia. These compensatory changes were evident in all task phases, even when no peg was grasped, indicating that the acute removal of tactile feedback is associated with non-specific modulation of eye-hand coordination.

We first determined whether digital anesthesia impacted performance on the pegboard task. Timing and force data revealed the importance of cutaneous responses to task performance. First, participants took more time to complete pick up and delivery following anesthesia, regardless of the precision required. Second, participants produced greater normal forces on the pegboard during all collection events and greater torque forces during precise collection events. Additionally, participants produced greater torque forces during precise delivery events. These results imply that somatosensory feedback is critical for force regulation and the torque force changes may reflect strategy shifts (e.g., hooking the peg) for precise collection and delivery events. Lastly, peg transport times were longer following anesthesia. Surprisingly, even movement of just the hand during coarse collection, without grasping a peg, required more time. These timing results suggest general adaptive changes that are not linked to specific task phases or even grasping. Collectively, these findings confirm that digital anesthesia impacted participants’ ability to perform the pegboard task. Although participants were able to complete the task, the duration and force changes indicate that adaptations following the loss of somatosensory function only provided a partial rescue of function.

Our main finding was that digital anesthesia resulted in reduced spatial separation between gaze and hand position. This change in gaze behavior was evident in all task phases. Following anesthesia, a smaller distance separated gaze and hand positions during pick up and collection events, which suggests that participants relied more on visual feedback for peg interactions following the loss of somatosensory signals. The most obvious effect is seen with coarse collection (Fig. 5C) where gaze is consistently close to the hand throughout the collection epoch in the anesthetized condition. In contrast, there is a greater eye-hand separation in all other conditions that increases over the coarse collection epoch as participants looked away from the pegs and toward the upcoming placement targets. Gaze changes were more subtle with precise actions, which may reflect a greater baseline demand of precise actions for visual support even with intact somatosensation. Indeed, gaze is closely tied to hand position during precise delivery and collection in all conditions, and anesthesia-related gaze changes are only evident at the end of the action epochs. This pattern reflects vision’s role in providing feedback regarding peg placement and collection success. However, with intact tactile sensation, vision is more quickly released from this feedback function, presumably to guide the upcoming hand movement, while vision persists in this function throughout the action epoch after anesthesia. These collective results support a conceptual framework in which vision and somatosensation serve as action-phase controllers during dexterous manipulation tasks that require eye-hand coordination (Johansson et al., 2001) and imply that vision becomes essential for achieving task subgoals when tactile ‘control points’ are unreliable.

We found that participants look closer to their hands during movement phases of the task following digital anesthesia. This effect was clearest with peg transport towards precise placement (Fig. 5D). Prior to anesthesia and in the control conditions, gaze and hand behaviors appear to be independent. Accordingly, the transport epoch is characterized by a large initial separation between gaze and hand positions as participants look toward the peg board targets while independently completing peg pick up. The eye-hand separation then closes over the transport epoch as the peg is transported to the pegboard target. We observed a different pattern with anesthesia: Gaze remained closely tied to the hand throughout the transport epoch. This pattern partially reflects the eye-hand relationship during the preceding (pick up) event, but it also reveals that participants looked closer to their hands as they transported the pegs, conceivably to monitor whether grasp was successfully maintained. This pattern is consistent with the hypothesis that vision is recruited to provide feedback when somatosensation is perturbed and we observed a similar pattern during transport for coarse delivery. Moreover, we found that during anesthesia participants looked closer to their hands and needed more time in movement phases when they were not grasping a peg. This surprising result implies that gaze behavior changes are not specifically tied to phases comprising object interactions, but also to the preparatory phase before object contact. This may represent compensation for loss of proprioceptive sensation in the anesthetized fingers and perturbed preshaping of the hand for peg grasping. Together these data support the notion that adaptation of eye-hand coordination following acute somatosensory feedback loss may be implemented through general mechanisms.

There are some study limitations to note. First, we have a relatively small sample size with only 5 participants. However, within the 5 participants we have a large sample of events given the number of sessions and trials performed by each participant. These data revealed high consistency within a subject, and individual differences that likely reflected the strategies employed to complete the task. We leveraged this variance in our GLMMs which included trial-level data from the experimental and control hands of each subject. In our study, 4 of the participants were right-hand dominant. Ideally, there would be an even split of right-hand dominant and left-hand dominant participants. Given the population distribution of right-hand vs. left-hand dominant participants, recruitment of left-handed participants is difficult. To partially control for this, we consistently used the dominant hand as the control hand and the non-dominant hand as the experimental hand. Of course, this choice also poses issues, namely that participants may have employed different strategies which using their dominant and non-dominant hands. We accounted for these differences by including hand dominance as a fixed effect in our GLMMs (i.e., whether the trial was completed using the dominant or non-dominant hand). Indeed, we observed variation in some force features explained by a main effect of hand; however, these changes were rarely observed in features where there was not also a significant hand X treatment interaction and the main effects were often smaller relative to the interaction effects. Another potential flaw with the experimental design could be the level of detail in the instructions. A defined strategy may have been more apparent with greater structure in the task; a specified order of precise delivery or collection would have allowed further analysis into the force feature outcomes and gaze behavior based on position. Lastly, our analyses of eye-hand relationships were handicapped by the limited sampling rate of our eye tracking system. With a higher sampling rate, we may have observed more sensitive dynamics in eye-hand coordination.

In conclusion, we found that the loss of somatosensory feedback through digital anesthesia impairs performance on a manual dexterity task. Behavioral adaptations following anesthesia only partially rescue performance. To compensate for the loss of touch, participants consistently directed their gaze closer to their hands, which we interpret as the recruitment of vision for feedback functions typically supported by touch. That gaze became more closely tied to the hands following anesthesia even in the absence of peg grasping implies that adaptations to dexterous behaviors maybe more general in nature. Whether adaptations also exist for prehension phases (object-free hand movement), during dexterous interactions with objects, and in action phase controllers signaling subgoal achievement in persons that suffer from chronic somatosensory feedback impairment could be studied using the compound eye-hand coordination assessment technique used here. Similarly, this approach provides biomarkers of adaptive changes after peripheral nerve injury affecting sensation in hand and the impact of clinical interventions. Finally, expanding our understanding of how dexterous behavior patterns change with altered somatosensory and visual feedback may be important for the use of neuroprosthetics while also informing training regimens for advanced robotic telesurgery.

## Extended Figure Legends

**Table 3-1.** Generalized linear mixed-effect model results for coarse collection. Model is fit to trials where all necessary data are available for the specific feature. Control hand, before anesthesia is taken as the baseline condition. Degrees of freedom (df), estimated using the Satterthwaite method, have been rounded to the nearest integer.

| Fixed Effects | Estimate | SE | df | t Value | P Value |
| --- | --- | --- | --- | --- | --- |
| Collection Tray Force Duration (ms) |  |  |  |  |  |
| Intercept | 347.98 | 57.25 | 5 | 6.078 | 0.00139 |
| *Intercept effects* |  |  |  |  |  |
| Treatment | -31.36 | 33.15 | 3008 | -0.946 | 0.34422 |
| Hand | 29.25 | 33.73 | 3002 | 0.867 | 0.38587 |
| Treatment:Hand | 783.82 | 46.95 | 3001 | 16.696 | < 2e-16 |
| Collection Tray Force Peak (N) |  |  |  |  |  |
| Intercept | 2.318 | 0.460 | 4 | 5.043 | 0.00707 |
| *Intercept effects* |  |  |  |  |  |
| Treatment | -0.173 | 0.053 | 3002 | -3.253 | 0.00115 |
| Hand | -0.051 | 0.054 | 3001 | -0.937 | 0.3489 |
| Treatment:Hand | 1.521 | 0.075 | 3000 | 20.162 | < 2e-16 |
| Collection Tray Torque Force (N*m) |  |  |  |  |  |
| Intercept | 9.346 | 1.505 | 4 | 6.21 | 0.0024 |
| *Intercept effects* |  |  |  |  |  |
| Treatment | 0.071 | 0.629 | 2464 | 0.114 | 0.9096 |
| Hand | -6.138 | 0.615 | 2463 | -9.978 | <2e-16 |
| Treatment:Hand | 1.363 | 0.890 | 2463 | 1.531 | 0.126 |

**Table 3-2.** Generalized linear mixed-effect model results for precise delivery. Model is fit to trials where all necessary data are available for the specific feature. Control hand, before anesthesia is taken as the baseline condition. Degrees of freedom (df), estimated using the Satterthwaite method, have been rounded to the nearest integer.

| Fixed Effects | Estimate | SE | df | t Value | P Value |
| --- | --- | --- | --- | --- | --- |
| Peg-hole Force Duration (ms) |  |  |  |  |  |
| Intercept | 260.5 | 38.26 | 10 | 6.807 | 3.73E-05 |
| *Intercept effects* |  |  |  |  |  |
| Treatment | 1.328 | 38.64 | 3143 | 0.034 | 0.97 |
| Hand | 23.91 | 38.74 | 3144 | 0.617 | 0.537 |
| Treatment:Hand | 573.8 | 54.92 | 3145 | 10.449 | < 2e-16 |
| Peg-hole Peak Force (N) |  |  |  |  |  |
| Intercept | 1.867 | 0.246 | 4 | 7.6 | 1.48E-03 |
| *Intercept effects* |  |  |  |  |  |
| Treatment | 0.389 | 0.040 | 3143 | 9.616 | < 2e-16 |
| Hand | 0.242 | 0.041 | 3144 | 5.986 | 2.40E-09 |
| Treatment:Hand | -1.118 | 0.057 | 3145 | -19.472 | < 2e-16 |
| Peg-hole Torque Force (N*m) |  |  |  |  |  |
| Intercept | 4.990 | 0.966 | 5 | 5.168 | 0.00335 |
| *Intercept effects* |  |  |  |  |  |
| Treatment | 1.638 | 0.560 | 1902 | 2.925 | 0.00348 |
| Hand | 1.096 | 0.553 | 1554 | 1.981 | 0.04772 |
| Treatment:Hand | 0.200 | 0.796 | 1776 | 0.251 | 0.80175 |

**Table 3-3.**
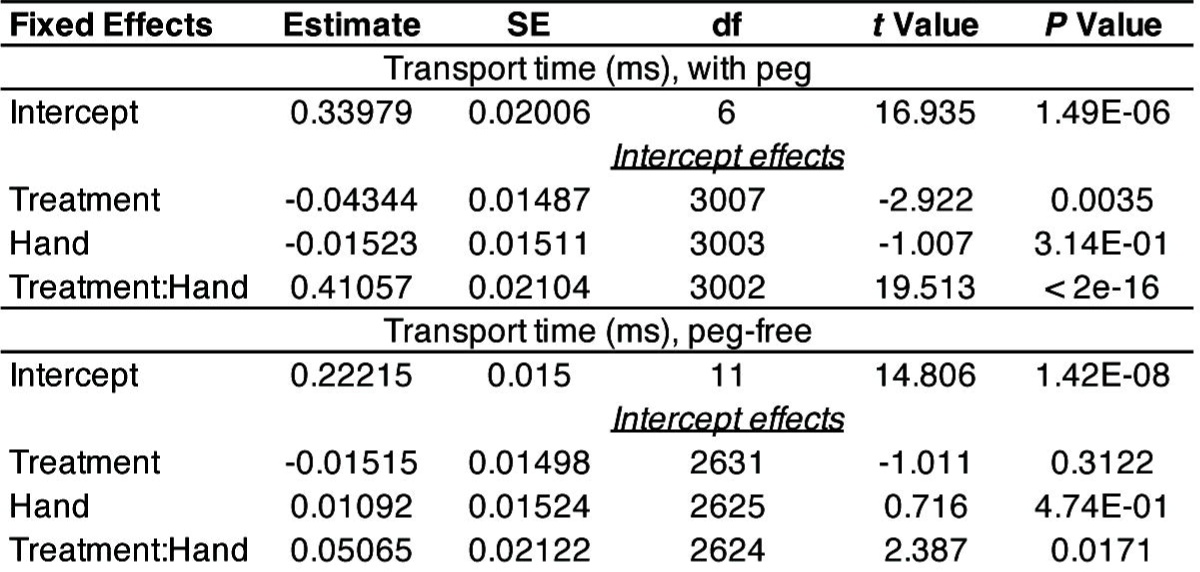
Generalized linear mixed-effect model results for transport time during placement phase. Model is fit to trials where all necessary data are available for the specific feature. Control hand, before anesthesia is taken as the baseline condition. Degrees of freedom (df), estimated using the Satterthwaite method, have been rounded to the nearest integer.

**Table 4-1.** Generalized linear mixed-effect model results for precise collection. Model is fit to trials where all necessary data are available for the specific feature. Control hand, before anesthesia is taken as the baseline condition. Degrees of freedom (df), estimated using the Satterthwaite method, have been rounded to the nearest integer.

| Fixed Effects | Estimate | SE | df | t Value | P Value |
| --- | --- | --- | --- | --- | --- |
| Peg-hole Force Duration (ms) |  |  |  |  |  |
| Intercept | 86.61 | 8.28 | 6 | 10.455 | 3.16E-05 |
| *Intercept effects* |  |  |  |  |  |
| Treatment | -5.99 | 6.14 | 3152 | -0.976 | 0.33 |
| Hand | 6.63 | 6.12 | 3151 | 1.084 | 0.28 |
| Treatment:Hand | 84.77 | 8.67 | 3152 | 9.78 | < 2e-16 |
| Peg-hole Peak Force (N) |  |  |  |  |  |
| Intercept | 0.576 | 0.108 | 4 | 5.319 | 5.57E-03 |
| *Intercept effects* |  |  |  |  |  |
| Treatment | 0.011 | 0.020 | 3152 | 0.535 | 0.59 |
| Hand | 0.048 | 0.020 | 3151 | 2.371 | 0.02 |
| Treatment:Hand | 0.208 | 0.029 | 3152 | 7.286 | 4.00E-13 |
| Peg-hole Torque Force (N*m) |  |  |  |  |  |
| Intercept | 1.305 | 0.280 | 5 | 4.654 | 0.004701 |
| *Intercept effects* |  |  |  |  |  |
| Treatment | 0.110 | 0.169 | 1934 | 0.652 | 0.514399 |
| Hand | 0.332 | 0.167 | 1612 | 1.994 | 4.63E-02 |
| Treatment:Hand | 0.835 | 0.240 | 1815 | 3.478 | 0.000517 |

**Table 4-2.** Generalized linear mixed-effect model results for coarse delivery. Model is fit to trials where all necessary data are available for the specific feature. Control hand, before anesthesia is taken as the baseline condition. Degrees of freedom (df), estimated using the Satterthwaite method, have been rounded to the nearest integer.

| Fixed Effects | Estimate | SE | df | t Value | P Value |
| --- | --- | --- | --- | --- | --- |
| Collection Tray Force Duration (ms) |  |  |  |  |  |
| Intercept | 85.12 | 5.56 | 5 | 15.3 | 1.96E-05 |
| *Intercept effects* |  |  |  |  |  |
| Treatment | -6.72 | 3.03 | 3153 | -2.217 | 0.027 |
| Hand | 11.96 | 3.02 | 3152 | 3.96 | 7.65E-05 |
| Treatment:Hand | 31.27 | 4.28 | 3153 | 7.313 | 3.29E-13 |
| Collection Tray Peak Force (N) |  |  |  |  |  |
| Intercept | 0.47 | 5.80E-02 | 4 | 8.081 | 0.001 |
| *Intercept effects* |  |  |  |  |  |
| Treatment | -0.08 | 1.57E-02 | 3152 | -4.876 | 1.14E-06 |
| Hand | 0.02 | 1.57E-02 | 3152 | 1.041 | 0.298 |
| Treatment:Hand | -0.04 | 2.22E-02 | 3152 | -1.955 | 0.051 |
| Collection Tray Torque Force (N*m) |  |  |  |  |  |
| Intercept | 0.594 | 0.093 | 5 | 6.413 | 0.00201 |
| *Intercept effects* |  |  |  |  |  |
| Treatment | -0.164 | 0.036 | 2502 | -4.54 | 5.90E-06 |
| Hand | -0.068 | 0.035 | 2601 | -1.943 | 0.052 |
| Treatment:Hand | 0.086 | 0.051 | 2476 | 1.683 | 0.09257 |

**Table 4-3.** Generalized linear mixed-effect model results for transport time during retrieval phase. Model is fit to trials where all necessary data are available for the specific feature. Control hand, before anesthesia is taken as the baseline condition. Degrees of freedom (df), estimated using the Satterthwaite method, have been rounded to the nearest integer.

| Fixed Effects | Estimate | SE | df | t Value | P Value |
| --- | --- | --- | --- | --- | --- |
| Transport time (ms), with peg |  |  |  |  |  |
| Intercept | 0.2326 | 0.01356 | 5.051 | 17.154 | 1.13E-05 |
| *Intercept effects* |  |  |  |  |  |
| Treatment | -0.03197 | 0.007344 | 3139 | -4.354 | 1.38E-05 |
| Hand | -0.004451 | 0.007305 | 3138 | -0.609 | 5.42E-01 |
| Treatment:Hand | 0.09718 | 0.01035 | 3138 | 9.394 | < 2e-16 |
| Transport time (ms), peg-free |  |  |  |  |  |
| Intercept | 0.205 | 0.016 | 5.24 | 13.055 | 3.41E-05 |
| *Intercept effects* |  |  |  |  |  |
| Treatment | -0.032 | 0.009 | 2835 | -3.505 | 4.63E-04 |
| Hand | -0.021 | 0.009 | 2835 | -2.36 | 1.83E-02 |
| Treatment:Hand | -0.037 | 0.013 | 2835 | -2.917 | 3.56E-03 |

**Table 5-1.**
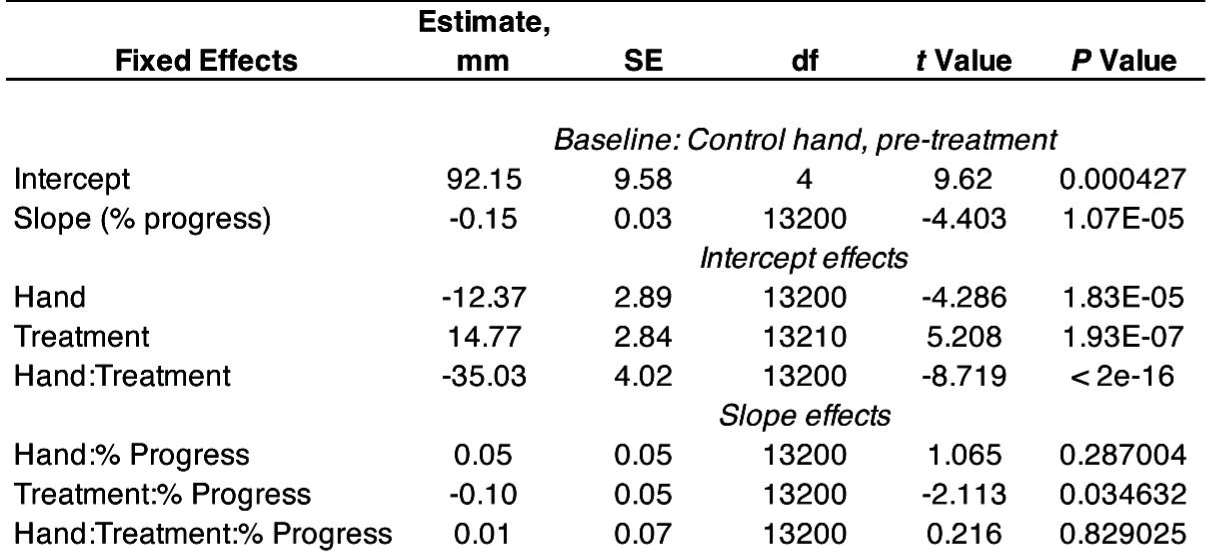
Generalized linear mixed-effect model results for GISI before coarse collection. Model is fit to trials where all necessary data are available for the specific feature. The first trial of each set was excluded because the hand begins at the starting position, rather than coming from the previous placement action. Control hand, before anesthesia is taken as the baseline condition. Degrees of freedom (df), estimated using the Satterthwaite method, have been rounded to the nearest integer.

**Table 5-2.** Generalized linear mixed-effect model results for GISI during peak force. Model is fit to trials where all necessary data are available for the specific feature. Control hand, before anesthesia is taken as the baseline condition. Degrees of freedom (df), estimated using the Satterthwaite method, have been rounded to the nearest integer.

| Fixed Effects | Estimate | SE | df | t Value | P Value |
| --- | --- | --- | --- | --- | --- |
| *Coarse collection (N)* |  |  |  |  |  |
| Intercept | 108.086 | 10.86 | 4 | 9.953 | 0.000392 |
| *Intercept effects* |  |  |  |  |  |
| Treatment | 12.379 | 3.384 | 3002 | 3.658 | 0.000258 |
| Hand | 14.794 | 3.326 | 3007 | 4.447 | 9.01E-06 |
| Treatment:Hand | -99.555 | 4.71 | 3001 | -21.137 | < 2e-16 |
| *Precise delivery (N)* |  |  |  |  |  |
| Intercept | 30.1619 | 5.4155 | 5 | 5.57 | 0.00341 |
| *Intercept effects* |  |  |  |  |  |
| Treatment | 3.4398 | 2.2475 | 3143 | 1.53 | 1.26E-01 |
| Hand | 0.1813 | 2.2417 | 3143 | 0.081 | 9.36E-01 |
| Treatment:Hand | 1.714 | 3.186 | 3144 | 0.538 | 5.91E-01 |
| *Precise collection (N)* |  |  |  |  |  |
| Intercept | 55.57 | 22.082 | 4 | 2.517 | 0.064626 |
| *Intercept effects* |  |  |  |  |  |
| Treatment | 10.59 | 3.202 | 3151 | 3.307 | 9.53E-04 |
| Hand | 4.053 | 3.211 | 3152 | 1.262 | 0.207057 |
| Treatment:Hand | -14.387 | 4.536 | 3152 | -3.172 | 1.53E-03 |
| *Coarse delivery (N)* |  |  |  |  |  |
| Intercept | 99.372 | 17.569 | 4 | 5.656 | 0.00449 |
| *Intercept effects* |  |  |  |  |  |
| Treatment | 1.396 | 3.059 | 3158 | 0.457 | 0.64802 |
| Hand | 6.406 | 3.067 | 3159 | 2.089 | 0.03679 |
| Treatment:Hand | -39.9 | 4.326 | 3158 | -9.223 | < 2e-16 |

**Table 5-3.** Generalized linear mixed-effect model results for GISI during coarse collection. Model is fit to trials where all necessary data are available for the specific feature. Control hand, before anesthesia is taken as the baseline condition. Degrees of freedom (df), estimated using the Satterthwaite method, have been rounded to the nearest integer.

| Fixed Effects | Estimate,<br>mm | SE | df | t Value | P Value |
| --- | --- | --- | --- | --- | --- |
| *Baseline: Control hand, pre-treatment* |  |  |  |  |  |
| Intercept | 71.01 | 13.41 | 4 | 5.294 | 0.00552 |
| Slope (% progress) | 0.48 | 0.03 | 15090 | 15.516 | < 2e-16 |
| *Intercept effects* |  |  |  |  |  |
| Hand | -12.35 | 2.66 | 15090 | -4.639 | 3.54E-06 |
| Treatment | 3.24 | 2.62 | 15100 | 1.238 | 0.21571 |
| Hand:Treatment | -28.23 | 3.71 | 15090 | -7.617 | 2.76E-14 |
| *Slope effects* |  |  |  |  |  |
| Hand:% Progress | 0.19 | 0.04 | 15090 | 4.271 | 1.96E-05 |
| Treatment:% Progress | 0.09 | 0.04 | 15090 | 2.203 | 0.02761 |
| Hand:Treatment:% Progress | -0.76 | 0.06 | 15090 | -12.6 | < 2e-16 |

**Table 5-4.**
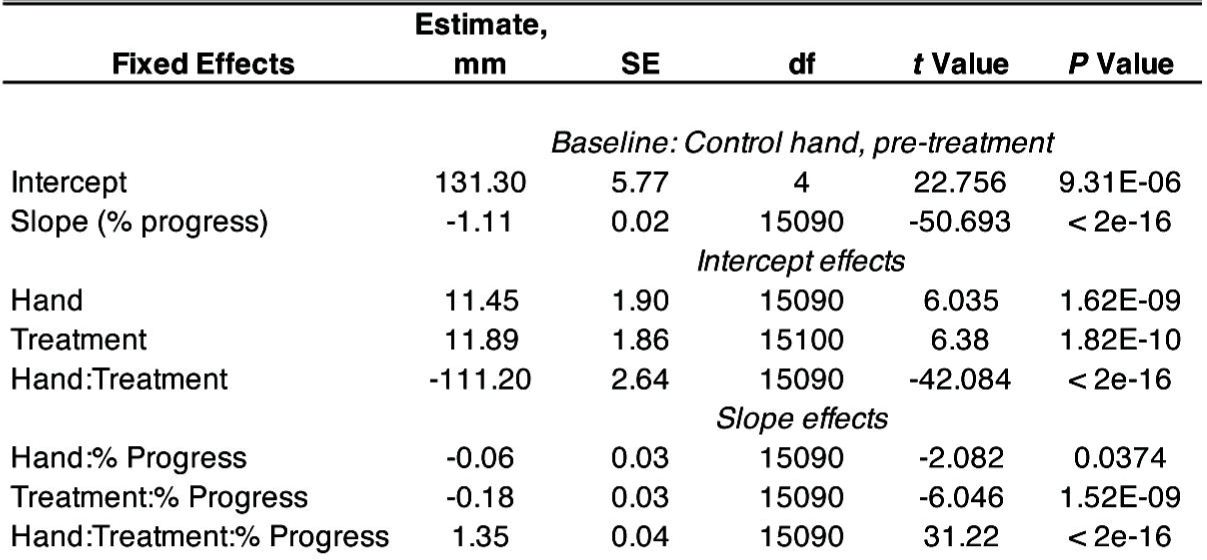
Generalized linear mixed-effect model results for GISI before precise delivery. Model is fit to trials where all necessary data are available for the specific feature. Control hand, before anesthesia is taken as the baseline condition. Degrees of freedom (df), estimated using the Satterthwaite method, have been rounded to the nearest integer.

**Table 5-5.** Generalized linear mixed-effect model results for GISI during precise delivery. Model is fit to trials where all necessary data are available for the specific feature. Control hand, before anesthesia is taken as the baseline condition. Degrees of freedom (df), estimated using the Satterthwaite method, have been rounded to the nearest integer.

| Fixed Effects | Estimate,<br>mm | SE | df | t Value | P Value |
| --- | --- | --- | --- | --- | --- |
| *Baseline: Control hand, pre-treatment* |  |  |  |  |  |
| Intercept | 16.61 | 6.12 | 4 | 2.713 | 0.048754 |
| Slope (% progress) | 0.35 | 0.02 | 15810 | 16.348 | < 2e-16 |
| *Intercept effects* |  |  |  |  |  |
| Hand | 5.77 | 1.84 | 15810 | 3.136 | 0.001713 |
| Treatment | -9.05 | 1.84 | 15810 | -4.931 | 8.27E-07 |
| Hand:Treatment | 18.95 | 2.61 | 15810 | 7.258 | 4.11E-13 |
| *Slope effects* |  |  |  |  |  |
| Hand:% Progress | -0.10 | 0.03 | 15810 | -3.425 | 0.000617 |
| Treatment:% Progress | 0.22 | 0.03 | 15810 | 7.246 | 4.5E-13 |
| Hand:Treatment:% Progress | -0.39 | 0.04 | 15810 | -9.211 | < 2e-16 |

**Table 6-1.**
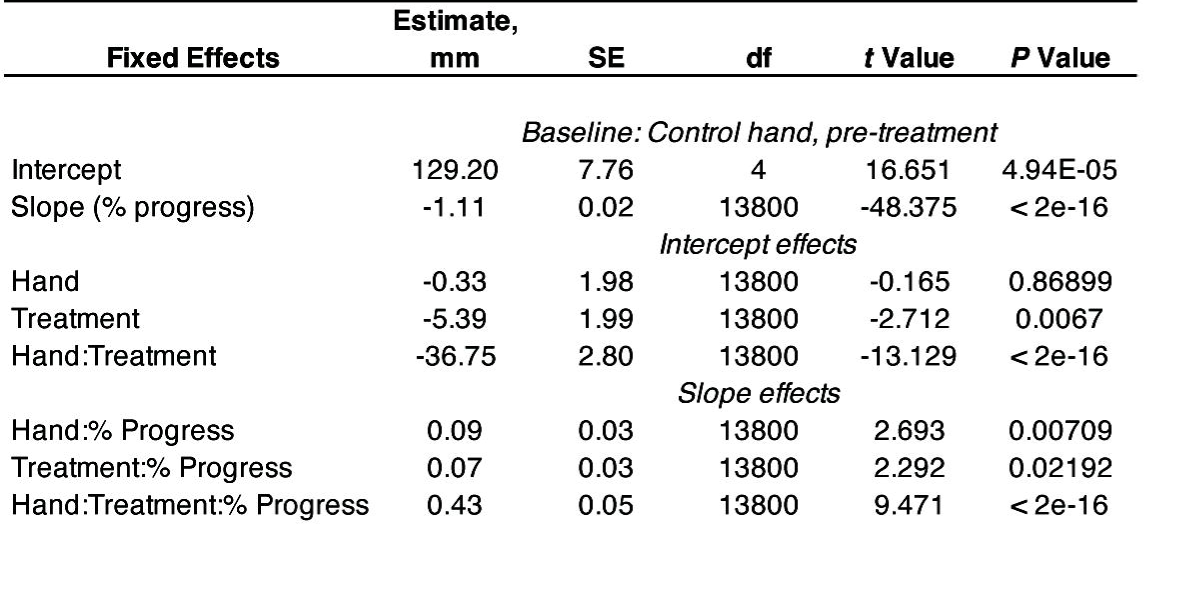
Generalized linear mixed-effect model results for GISI before precise collection. Model is fit to trials where all necessary data are available for the specific feature. The first trial of each set was excluded because the hand begins at the starting position, rather than coming from the previous placement action. Control hand, before anesthesia is taken as the baseline condition. Degrees of freedom (df), estimated using the Satterthwaite method, have been rounded to the nearest integer.

**Table 6-2.** Generalized linear mixed-effect model results for GISI during precise collection. Model is fit to trials where all necessary data are available for the specific feature. Control hand, before anesthesia is taken as the baseline condition. Degrees of freedom (df), estimated using the Satterthwaite method, have been rounded to the nearest integer.

| Fixed Effects | Estimate,<br>mm | SE | df | t Value | P Value |
| --- | --- | --- | --- | --- | --- |
| *Baseline: Control hand, pre-treatment* |  |  |  |  |  |
| Intercept | 24.37 | 19.34 | 4 | 1.26 | 0.2752 |
| Slope (% progress) | 0.58 | 0.03 | 15850 | 21.023 | < 2e-16 |
| *Intercept effects* |  |  |  |  |  |
| Hand | 13.61 | 2.38 | 15850 | 5.711 | 1.14E-08 |
| Treatment | -3.01 | 2.39 | 15850 | -1.258 | 0.2084 |
| Hand:Treatment | 2.84 | 3.38 | 15850 | 0.843 | 0.3994 |
| *Slope effects* |  |  |  |  |  |
| Hand:% Progress | -0.09 | 0.04 | 15850 | -2.318 | 0.0205 |
| Treatment:% Progress | 0.08 | 0.04 | 15850 | 2.018 | 0.0436 |
| Hand:Treatment:% Progress | -0.25 | 0.06 | 15850 | -4.599 | 4.27E-06 |

**Table 6-3.** Generalized linear mixed-effect model results for GISI before coarse delivery. Model is fit to trials where all necessary data are available for the specific feature. Control hand, before anesthesia is taken as the baseline condition. Degrees of freedom (df), estimated using the Satterthwaite method, have been rounded to the nearest integer.

| Fixed Effects | Estimate,<br>Pixels | SE | df | t Value | P Value |
| --- | --- | --- | --- | --- | --- |
| *Baseline: Control hand, pre-treatment* |  |  |  |  |  |
| Intercept | 99.32 | 11.35 | 4 | 8.749 | 0.000735 |
| Slope (% progress) | -0.20 | 0.03 | 15770 | -6.222 | 5.03E-10 |
| *Intercept effects* |  |  |  |  |  |
| Hand | -5.02 | 2.72 | 15770 | -1.847 | 0.064796 |
| Treatment | -1.42 | 2.73 | 15770 | -0.52 | 0.602764 |
| Hand:Treatment | -21.90 | 3.85 | 15770 | -5.686 | 1.33E-08 |
| *Slope effects* |  |  |  |  |  |
| Hand:% Progress | 0.13 | 0.04 | 15770 | 2.87 | 0.004112 |
| Treatment:% Progress | 0.16 | 0.04 | 15770 | 3.594 | 0.000326 |
| Hand:Treatment:% Progress | -0.10 | 0.06 | 15770 | -1.656 | 0.097837 |

**Table 6-4.** Generalized linear mixed-effect model results for GISI during coarse delivery. Model is fit to trials where all necessary data are available for the specific feature. Control hand, before anesthesia is taken as the baseline condition. Degrees of freedom (df), estimated using the Satterthwaite method, have been rounded to the nearest integer.

| Fixed Effects | Estimate,<br>mm | SE | df | t Value | P Value |
| --- | --- | --- | --- | --- | --- |
| *Baseline: Control hand, pre-treatment* |  |  |  |  |  |
| Intercept | 80.99 | 18.05 | 4 | 4.488 | 0.0105 |
| Slope (% progress) | 0.32 | 0.03 | 15880 | 11.83 | < 2e-16 |
| *Intercept effects* |  |  |  |  |  |
| Hand | 9.84 | 2.37 | 15880 | 4.157 | 3.24E-05 |
| Treatment | 16.55 | 2.37 | 15880 | 6.973 | 3.22E-12 |
| Hand:Treatment | -46.29 | 3.35 | 15880 | -13.823 | < 2e-16 |
| *Slope effects* |  |  |  |  |  |
| Hand:% Progress | -0.06 | 0.04 | 15880 | -1.527 | 0.1268 |
| Treatment:% Progress | -0.19 | 0.04 | 15880 | -4.944 | 7.73E-07 |
| Hand:Treatment:% Progress | 0.05 | 0.05 | 15880 | 0.94 | 0.3474 |

## References

Bilaloglu, S., Lu, Y., Geller, D., Ross Rizzo, J., Aluru, V., Gardner, E. P., & Raghavan, P. (2016). Effect of blocking tactile information from the fingertips on adaptation and execution of grip forces to friction at the grasping surface. J Neurophysiol, 115, 1122–1131. 10.1152/jn.00639.2015.-Adaptation

Birznieks, I., Burstedt, M. K. O., Edin, B. B., & Johansson, R. S. (1998). Mechanisms for Force Adjustments to Unpredictable Frictional Changes at Individual Digits During Two-Fingered Manipulation. Journal of Neurophysiology, 80(4), 1989–2002. 10.1152/jn.1998.80.4.1989

Caplan, B., & Mendoza, J. E. (2011). Edinburgh Handedness Inventory. In Encyclopedia of Clinical Neuropsychology (pp. 928–928). Springer New York. 10.1007/978-0-387-79948-3_684

Chesler, A. T., Szczot, M., Bharucha-Goebel, D., Čeko, M., Donkervoort, S., Laubacher, C., Hayes, L. H., Alter, K., Zampieri, C., Stanley, C., Innes, A. M., Mah, J. K., Grosmann, C. M., Bradley, N., Nguyen, D., Foley, A. R., Le Pichon, C. E., & Bönnemann, C. G. (2016). The Role of *PIEZO2* in Human Mechanosensation. New England Journal of Medicine, 375(14), 1355–1364. 10.1056/NEJMoa1602812

Gordon, A. M., Westling, G., Cole, K. J., & Johansson, R. S. (1993). Memory representations underlying motor commands used during manipulation of common and novel objects. Journal of Neurophysiology, 69(6), 1789–1796. 10.1152/jn.1993.69.6.1789

Jenmalm, P., Birznieks, I., Goodwin, A. W., & Johansson, R. S. (2003). Influence of object shape on responses of human tactile afferents under conditions characteristic of manipulation. European Journal of Neuroscience, 18(1), 164–176. 10.1046/j.1460-9568.2003.02721.x

Jenmalm, P., Dahlstedt, S., & Johansson, R. S. (2000). Visual and Tactile Information About Object-Curvature Control Fingertip Forces and Grasp Kinematics in Human Dexterous Manipulation. Journal of Neurophysiology, 84(6), 2984–2997. 10.1152/jn.2000.84.6.2984

Jenmalm, P., & Johansson, R. S. (1997). Visual and Somatosensory Information about Object Shape Control Manipulative Fingertip Forces. The Journal of Neuroscience, 17(11), 4486–4499. 10.1523/JNEUROSCI.17-11-04486.1997

Johansson, R. S., & Flanagan, J. R. (2009). Coding and use of tactile signals from the fingertips in object manipulation tasks. In Nature Reviews Neuroscience (Vol. 10, Issue 5, pp. 345–359). 10.1038/nrn2621

Johansson, R. S., & Westling, G. (1984). Roles of glabrous skin receptors and sensorimotor memory in automatic control of precision grip when lifting rougher or more slippery objects. Experimental Brain Research, 56(3). 10.1007/BF00237997

Johansson, R. S., & Westling, G. (1987). Signals in tactile afferents from the fingers eliciting adaptive motor responses during precision grip. Experimental Brain Research, 66(1). 10.1007/BF00236210

Johansson, R. S., Westling, G., Bäckström, A., & Flanagan, J. R. (2001). Eye–Hand Coordination in Object Manipulation. The Journal of Neuroscience, 21(17), 6917–6932. 10.1523/JNEUROSCI.21-17-06917.2001

Johnson, K. O., & Hsiao, S. S. (1992). Neural Mechanisms of Tactual form and Texture Perception. Annual Review of Neuroscience, 15(1), 227–250. 10.1146/annurev.ne.15.030192.001303

Killick, R., Fearnhead, P., & Eckley, I. A. (2012). Optimal Detection of Changepoints With a Linear Computational Cost. Journal of the American Statistical Association, 107(500), 1590–1598. 10.1080/01621459.2012.737745

Kuznetsova, A., Brockhoff, P. B., & Christensen, R. H. B. (2017). lmerTest Package: Tests in Linear Mixed Effects Models. Journal of Statistical Software, 82(13). 10.18637/jss.v082.i13

Lukos, J., Ansuini, C., & Santello, M. (2007). Choice of Contact Points during Multidigit Grasping: Effect of Predictability of Object Center of Mass Location. The Journal of Neuroscience, 27(14), 3894–3903. 10.1523/JNEUROSCI.4693-06.2007

Mahmud, A. A., Nahid, N. A., Nassif, C., Sayeed, M. S. B., Ahmed, M. U., Parveen, M., Khalil, M. I., Islam, M. M., Nahar, Z., Rypens, F., Hamdan, F. F., Rouleau, G. A., Hasnat, A., & Michaud, J. L. (2017). Loss of the proprioception and touch sensation channel <scp>PIEZO2</scp> in siblings with a progressive form of contractures. Clinical Genetics, 91(3), 470–475. 10.1111/cge.12850

Monzée, J., Lamarre, Y., & Smith, A. M. (2003). The effects of digital anesthesia on force control using a precision grip. Journal of Neurophysiology, 89(2), 672–683. 10.1152/jn.00434.2001

Nordmark, P. F., Ljungberg, C., & Johansson, R. S. (2018). Structural changes in hand related cortical areas after median nerve injury and repair. Scientific Reports, 8(1), 4485. 10.1038/s41598-018-22792-x

Rao, A., & Gordon, A. (2001). Contribution of tactile information to accuracy in pointing movements. Experimental Brain Research, 138(4), 438–445. 10.1007/s002210100717

Sailer, U., Flanagan, J. R., & Johansson, R. S. (2005). Eye–Hand Coordination during Learning of a Novel Visuomotor Task. The Journal of Neuroscience, 25(39), 8833–8842. 10.1523/JNEUROSCI.2658-05.2005

Sathian, K., Goodwin, A., John, K., & Darian-Smith, I. (1989). Perceived roughness of a grating: correlation with responses of mechanoreceptive afferents innervating the monkey’s fingerpad. The Journal of Neuroscience, 9(4), 1273–1279. 10.1523/JNEUROSCI.09-04-01273.1989

Sober, S. J., & Sabes, P. N. (2005). Flexible strategies for sensory integration during motor planning. Nature Neuroscience, 8(4), 490–497. 10.1038/nn1427

Westling, G., & Johansson, R. S. (1987). Responses in glabrous skin mechanoreceptors during precision grip in humans. Experimental Brain Research, 66(1). 10.1007/BF00236209

Winges, S. A., Weber, D. J., & Santello, M. (2003). The role of vision on hand preshaping during reach to grasp. Experimental Brain Research, 152(4), 489–498. 10.1007/s00221-003-1571-9

Yahya, A., Kluding, P., Pasnoor, M., Wick, J., Liu, W., & dos Santos, M. (2019). The impact of diabetic peripheral neuropathy on pinch proprioception. Experimental Brain Research, 237(12), 3165–3174. 10.1007/s00221-019-05663-3

